# Mitochondrial proline catabolism activates Ras1/cAMP/PKA-induced filamentation in *Candida albicans*

**DOI:** 10.1101/374777

**Authors:** Fitz Gerald S. Silao, Meliza Ward, Kicki Ryman, Axel Wallström, Björn Brindefalk, Klas Udekwu, Per O. Ljungdahl

## Abstract

Amino acids are among the earliest identified inducers of yeast-to-hyphal transitions in *Candida albicans*, an opportunistic fungal pathogen of humans. Here, we show that the morphogenic amino acids arginine, ornithine and proline are internalized and metabolized in mitochondria via a *PUT1*- and *PUT2*-dependent pathway that results in enhanced ATP production. Elevated ATP levels correlate with Ras1/cAMP/PKA pathway activation and Efg1-induced gene expression. The magnitude of amino acid-induced filamentation is linked to glucose availability; high levels of glucose repress mitochondrial function thereby dampening filamentation. Furthermore, arginine-induced morphogenesis occurs more rapidly and independently of Dur1,2-catalyzed urea degradation, indicating that mitochondrial-generated ATP, not CO_2_, is the primary morphogenic signal derived from arginine metabolism. The important role of the SPS-sensor of extracellular amino acids in morphogenesis is the consequence of induced amino acid permease gene expression, i.e., SPS-sensor activation enhances the capacity of cells to take up morphogenic amino acids, a requisite for their catabolism. *C. albicans* cells engulfed by murine macrophages filament, resulting in macrophage lysis. Phagocytosed *put1-/-* and *put2*-/- cells do not filament and do not lyse macrophages, consistent with a critical role of mitochondrial proline metabolism in virulence.

## Introduction

*Candida albicans* is an opportunistic fungal pathogen that commonly exists as a benign member of the human microbiome. Immunosuppression, or microbial dysbiosis, can predispose an individual to infection, enabling this fungus to initiate and develop a spectrum of pathologies, including superficial mucocutaneous or even life-threatening invasive infections [1, 2]. As a human commensal, *C. albicans* can asymptomatically colonize virtually all anatomical sites in the host, each with a characteristic and unique microenvironment, with differing nutrient and microbiome compositions, physical properties, and levels of innate immune defenses [3]. The ability to colonize and infect discrete microenvironments is attributed to an array of virulence characteristics, a major one being its morphological plasticity. As a pleomorphic organism, *C. albicans* can assume at least three distinct morphologies: yeast-like, pseudohyphae, and true hyphae, where the latter two are commonly referred to as filamentous morphologies (for review see [4–7]. Strains that are genetically locked in either yeast or filamentous forms fail to mount infections *in vitro* and *in vivo* infection models, supporting the concept that morphological switching, rather than the specific morphology *per se*, is a requisite to virulence [4, 6, 8–10]. The environmental signals known to trigger morphogenesis in *C. albicans* reflect the conditions within the human host, such as temperature (37 °C) and CO_2_, alkaline pH, the presence of serum, N-acetylglucosamine, and a discrete set of amino acids.

Early studies examining amino acid-induced morphogenesis implicated metabolism as being important for filamentation, and the inducing effects were shown to correlate to their specific point-of-entry in metabolism [11–13]. The most potent inducers of filamentation are amino acids that are catabolized to glutamate, such as arginine and proline, which enters the TCA cycle via α-ketoglutarate. Importantly, arginine and proline can supply nitrogen and carbon for intermediary metabolism and their catabolism provides energy to support diverse cellular functions. Studies examining proline uptake and distribution during filamentous growth suggested that proline catabolism results in an increase in the cellular reducing potential, i.e., enhanced levels of reduced flavoproteins were noted [11]. Several of the conclusions from these earlier studies, in particular that filamentous growth of *C. albicans* is linked to repression of mitochondrial activity [11–13], appear to conflict with more recent reports showing that filamentation is accompanied by increased mitochondrial respiratory activity [14–16]. Clearly, the underlying mechanisms through which amino acids induce filamentation remain to be defined. In particular, the basis of arginine- and proline-induced morphogenesis needs to be placed in context to the current mechanistic understanding of the signaling cascades implicated in morphogenesis.

Among the central metabolic signaling pathways in *C. albicans* linked to morphogenesis, the best characterized are the mitogen-activated protein kinase (MAPK) and the 3’-5’-cyclic adenosine monophosphate/Protein Kinase A (cAMP/PKA) signaling systems, which activate the transcription factors Cph1 and Efg1, respectively [8, 17, 18], reviewed in [4, 7, 19, 20]. Ras1 is a small GTPase required for proper MAPK and cAMP/PKA signaling, and specifically for the induction of filamentation by amino acids and serum [21, 22], reviewed in [20, 23]. Recently, Grahl et al. have proposed that intracellular ATP levels and increased mitochondrial activity, control the activation of Ras1/cAMP/PKA pathway [14]. In this intriguing model, the adenyl cyclase (Cyr1/Cdc35) works cooperatively in a positive feedback loop with ATP as key input. Accordingly, ATP promotes Cyr1 binding to the active GTP-bound form of Ras1 thereby reducing the ability of Ira2 to stimulate the intrinsic GTPase activity of Ras1. As a consequence, enhanced Cyr1 activity leads to elevated levels of cAMP and amplification of PKA-dependent signaling, activating the effector transcription factor Efg1 and the expression of genes required for filamentous growth [24–26], reviewed in [20, 23, 27, 28].

Some morphogenic signals appear to bypass the requirement for Ras1 (reviewed in [27, 28]. By example, CO_2_ is a well-characterized stimulus for morphological switching in *C. albicans*; CO_2_ binds directly and activates Cyr1 [29]. Ghosh et al. have proposed that arginine-induced morphogenesis is the consequence of arginase (*CAR1)* dependent metabolism to ornithine and urea, and subsequent urea amidolyase (*DUR1,2*) dependent generation of CO_2_ from urea [30]. Also, the G protein-coupled receptor Gpr1, which has been implicated in amino acid-induced morphogenesis, does not appear to require Ras1. Gpr1-initiated signals activate Cyr1 by stimulating GTP-GDP exchange on the Gα protein Gpa2; the active GTP-bound form of Gpa2 is thought to bind to the Gα-binding domain within the N-terminal of Cyr1 leading to enhanced cAMP production (reviewed in [20, 28]. It has been reported that Gpr1 senses the presence of extracellular methionine [31] and glucose [32], however recently, lactate has been proposed to be the primary activating ligand [33]. The role of Gpr1 in amino acid-induced morphogenesis remains an open question.

*C. albicans* cells respond to the presence of extracellular amino acids using the plasma membrane-localized SPS (Ssy1-Ptr3-Ssy5) sensor complex [34–36]. In response to amino acids, the primary sensor Ssy1 (Csy1) is stabilized in a signaling conformation leading to Ssy5-mediated proteolytic processing of two latently expressed transcription factors, Stp1 and Stp2 [34]. The processed factors efficiently target to the nucleus activating the expression of distinct sets of genes required for assimilation of external nitrogen. Stp1 regulates the expression of *SAP2*, encoding the major secreted aspartyl proteinase, and oligopeptide transporters (*OPT1* and *OPT3*); whereas Stp2, derepresses the expression of a subset of amino acid permeases (AAP) that facilitate amino acid uptake. *STP1* expression is controlled by nitrogen catabolite repression (NCR), a supra-regulatory system that represses that utilization of non-preferred nitrogen sources when preferred ones are available [37]. The endoplasmic reticulum (ER)-localized chaperone Csh3, is required for the functional expression of both Ssy1 and AAPs, and thus acts as the most upstream and downstream component of the SPS sensing pathway [35]. Strains lacking either Ssy1 or Csh3 fail to efficiently respond to the presence of extracellular amino acids and serum and exhibit impaired morphological switching [35, 36]. It has not previously been determined if the SPS-sensor induces morphogenesis directly in response to extracellular amino acids, or indirectly, the consequence of enhanced amino acid uptake and subsequent intracellular signaling events.

In this report, we show how amino acid-induced and SPS-sensor-dependent signals are integrated into the central signaling pathways that control yeast-to-hyphal morphological transitions in *C. albicans*. Our results indicate that the augmented levels of intracellular ATP, resulting from catabolism of proline in the mitochondria, correlate with activated Ras1/cAMP/PKA and Efg1-dependent gene expression. The magnitude of the response is sensitive to the levels of glucose in a manner consistent with glucose repression of mitochondrial function. The SPS-sensor plays an indirect, but important, role in enhancing the uptake of the inducing amino acids. Finally, we show that *C. albicans* cells express proline catabolic enzymes when phagocytosed by murine macrophages, and that inactivation of proline catabolism diminishes the capacity of *C. albicans* cells to induce hyphal growth and escape engulfing macrophages.

## Results

### Amino acid-induced morphogenesis is dependent on uptake

We assessed the capacity of ornithine, citrulline, and the 20 amino acids commonly found in proteins to induce filamentous growth in *C. albicans.* Wildtype (WT) cells were grown as macrocolonies on MES-buffered (pH of 6.0) synthetic dextrose (2% glucose) medium containing 10 mM of each individual amino acid as sole nitrogen source. As shown in **Fig. 1A**, proline and arginine strongly induced filamentous growth as evidenced by the formation of wrinkled macrocolonies. Microscopic evaluation of cells from wrinkled colonies confirmed the presence of extensive filamentous growth (mainly hyphae). Ornithine, a non-proteinogenic amino acid and a catabolic intermediate in the degradation of arginine, induced pronounced filamentous growth. Of the amino acids tested, aspartate consistently produced smooth macrocolonies comprised of round cells, exclusively yeast-like in appearance. Consequently, aspartate was chosen as a reference for subsequent studies.

**Fig 1.**
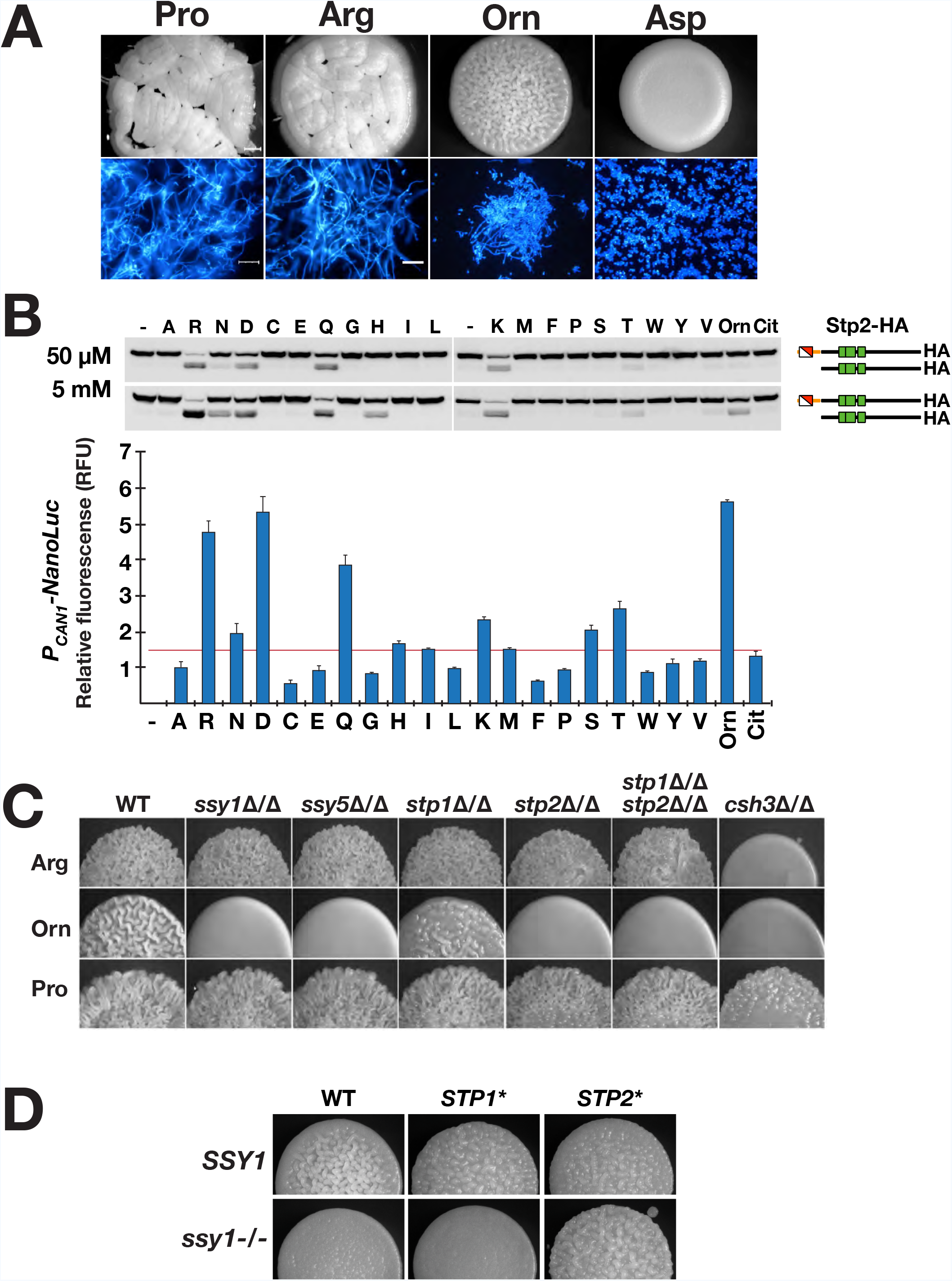
Amino acid-induced morphogenesis is dependent on amino acid uptake. **A.** Macrocolonies of wildtype *C. albicans* (PMRCA18) grown on S<u>X</u>D medium containing 10 mM of the indicated amino acids (<u>X</u> = Pro, Arg, Orn or Asp) and 2% glucose after 48 h of growth at 37 °C (upper panels). Cells scraped from macrocolonies stained with calcofluor white (lower panels); scale bars = 30 μ. **B.** Amino acid-induced SPS-sensor signaling. Cells expressing Stp2-6XHA (PMRCA48) were grown to log phase in SD medium and induced with 50 μM or 5 mM of the indicated amino acids for 5 min at 30 °C. The levels of latent and processed Stp2 in extracts were analyzed by immunoblotting (upper panels). Similarly, reporter strain (CFG001) carrying an integrated P_*CAN1*_-NanoLuc™-PEST construct was grown to log phase in SD medium and induced with 50 μM of the indicated amino acids for 2 h at 30 °C. The average luciferase signal (ave. ± CI, 95% CL) are plotted; threshold for significance ≥ 1.5X fold change). **C.** Macrocolonies of wildtype (WT; PMRCA18) and strains carrying mutations inactivating SPS-sensing pathway components *ssy1*Δ/Δ (YJA64), *ssy5*Δ/Δ (YJA53), *stp1*Δ/Δ (PMRCA59), *stp2*Δ/Δ (PMRCA57), *stp1*Δ/Δ *stp2*Δ/Δ (PMRCA94) and *csh3*Δ/Δ (PMRCA12) grown on the indicated S<u>X</u>D media. **D.** Constitutively active Stp2* but not Stp1* bypasses the filamentous growth defect of a *ssy1* null mutant in the presence of ornithine. Macrocolonies of WT (PMRCA18), *STP1\** (PMRCA23), *STP2\** (PMRCA44), *ssy1*-/- *STP1\** (CFG078), and *ssy1*-/-*STP2\** (CFG073) grown on S<u>O</u>D with ornithine (O) as sole nitrogen source. Images in C and D were obtained after 24 h of incubation 37 °C.

Using quantitative RT-PCR (qRT-PCR) we analyzed the expression of known hyphae-specific genes (HSG) *ECE1*, *EED1*, *HWP1*, *UME6*, *ALS3*, *HGC1*, *SAP4*, and *SAP5* [4] in cells from colonies grown on media with arginine, proline, ornithine and aspartate. With the exception of *EED1*, the expression of HSG were clearly induced in cells grown on media with morphogenic amino acids, ≥ 7-fold higher than in cells grown on aspartate **(Fig. S1)**. *SAP4,* a known Efg1-regulated gene [38], exhibited the highest level of induction, ≥ 80-fold higher levels than in aspartate grown cells. These experiments were repeated using liquid cultures, and the same trends were observed (data not shown). These results confirm the appropriateness of using macrocolonies to score amino acid-induced morphogenesis.

Next, we evaluated whether SPS-sensor activation was required for amino acid-induced morphogenesis. This was accomplished by assessing SPS-sensor dependent Stp2 processing [34]. A strain carrying a functional C-terminal HA tagged Stp2 (Stp2-HA; PMRCA44) was grown in minimal ammonium-based synthetic dextrose (SD) medium, extracts were prepared 5 min after induction by the indicated amino acid. Arginine (R), asparagine (N), aspartate (D), glutamine (Q), histidine (H), lysine (K), serine (S) and ornithine (Orn) efficiently activated the SPS-sensor; extracts contained the shorter processed form of Stp2 (**Fig. 1B**, upper panel). Next, we assessed Stp2-dependent promoter activation using an integrated P_*CAN1*_-NanoLuc™-PEST reporter construct; the expression of the luciferase signal is controlled by the *CAN1* promoter, which is strictly dependent on the SPS-sensor and Stp2 (**Fig. S2A and S2B**). The inclusion of the 41-amino acid PEST sequence confers a shorter NanoLuc^TM^ lifetime, which facilitates a tighter coupling of transcription and translation [39]. Enhanced luminescence was observed only in cells induced with the amino acids giving rise to Stp2 processing (**Fig. 1B**, lower panel). Notably, proline, which induces robust filamentation, did not activate the SPS-sensor as no Stp2 processing or luminescence was detected. Conversely, aspartate, which does not induce filamentous growth, robustly activated the SPS-sensor as determined by Stp2 processing and enhanced luciferase activity. These results indicate that amino acid-induced morphogenesis is not obligatorily coupled to SPS-sensor signaling.

The contribution of signals derived from the SPS sensing pathway on filamentation induced by arginine, ornithine and proline was examined. Arginine, a potent inducer of the SPS-sensor (**Fig. 1B**), induced filamentation in an SPS-sensor independent manner; filamentation was observed in mutants lacking components of the SPS sensing pathway (**Fig. 1C**). By contrast, ornithine, also a potent inducer of the SPS-sensor, induced filamentous growth in a strictly SPS-sensor- and Stp2-dependent manner (**Fig. 1C**). Notably, Stp1, a transcription factor that induces genes required for extracellular protein utilization, is not required for ornithine-induced filamentation. Proline, which does not induce SPS-sensor signaling, promoted filamentous growth in an SPS-sensor independent manner (**Fig. 1C**). Importantly, the filamentation was greatly reduced in cells lacking *CSH3* (*csh3*Δ/Δ), a gene encoding a membrane-localized chaperone required for the functional expression of Ssy1 and most amino acid permeases [35, 36] (**Fig. 1C**), clearly suggesting that amino acid uptake is required for amino acid-induced morphogenesis.

The clear requirement of the SPS-sensor in facilitating ornithine-induced filamentation provided the opportunity to rigorously test the notion that uptake is essential. Based on the knowledge that amino acid permease-dependent uptake is dependent on Stp2 and not Stp1, we used the CRISPR/Cas9 system to introduce *ssy1* null mutations in strains expressing constitutively active Stp2 (*STP2*\*) or Stp1 (*STP1*\*) (**Fig. S3A**). The results clearly show that *STP2*\*, but not *STP1*\* (**Fig. 1D**), bypasses the *ssy1* null mutation, indicating that the permease responsible for ornithine uptake is indeed encoded by a SPS-sensor and Stp2 controlled gene. Similarly, SPS-sensor dependence was observed for the inducing amino acids alanine, glutamine, and serine (data not shown). Together, these results indicate that amino acid-induced filamentous growth is dependent on the uptake of the inducing amino acid.

### Amino acid-induced morphogenesis is dependent on catabolism and Ras1/cAMP/PKA signaling

Two core signaling pathways, i.e., MAPK and cAMP/PKA, are known to transduce metabolic signals that affect filamentous growth (**Fig. 2A**). We evaluated the capacity of amino acids to induce filamentation in cells carrying null alleles of *RAS1* and the effector transcription factors, *CPH1* and *EFG1* diagnostic for MAPK and cAMP/PKA signaling, respectively [8, 18, 40](**Fig. 2B**). Similar to wildtype, *cph1*Δ/Δ cells were wrinkled in appearance, indicating that amino acid induced filamentation was independent of MAPK signaling. By contrast, the colonies derived from *ras1*Δ/Δ and *efg1*Δ/Δ cells were smooth. As expected, the *efg1*Δ/Δ *cph1*Δ/Δ double mutant strain also formed smooth colonies. These results indicate that the inducing signals are transduced by the cAMP/PKA pathway. A clear dependence on Ras1/cAMP/PKA signaling was also observed for other inducing amino acids, i.e., alanine, glutamine, and serine (data not shown). Our results demonstrating that amino acid-induced morphogenesis is strictly dependent on Ras1 is contrary to current models that postulate that amino acids-initiated signals are transduced by Gpr1/Gpa2 (reviewed in [20, 28]. According to these models, amino acid signaling should be Ras1 independent.

**Fig 2.**
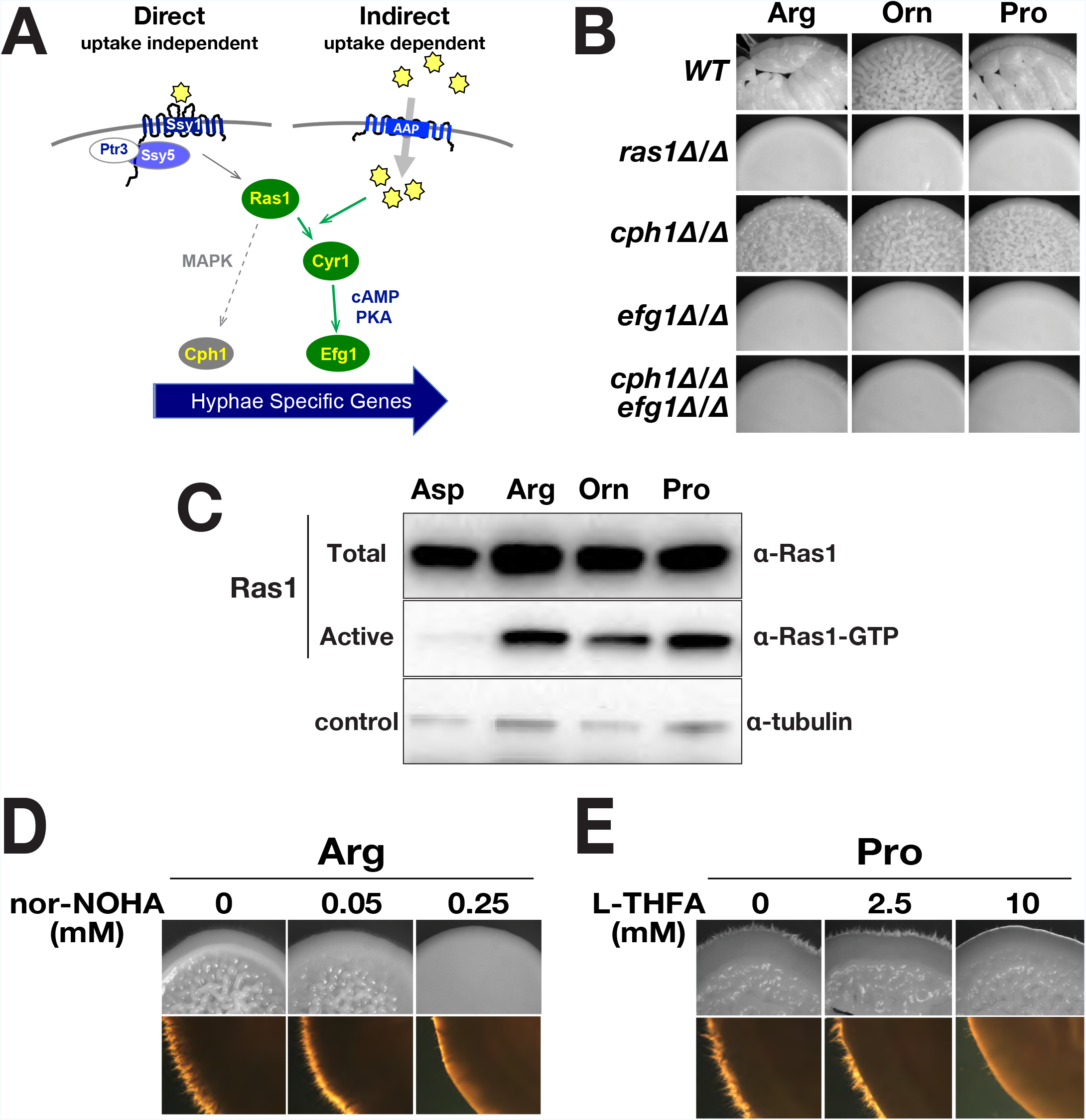
Amino acid-induced morphogenesis is dependent on catabolism and Ras1 activated Efg1-dependent transcription. **A.** Scheme of possible signaling pathways controlling amino acid-induced morphogenesis. **B.** Amino acid-induced morphogenesis requires a functional Ras1/cAMP/PKA pathway (Efg1-dependent) but not on the MAPK signaling pathway (Cph1-dependent). Wildtype (WT; PMRCA18) and strains lacking Ras1 (CDH107), Cph1 (JKC19), Efg1 (HLC52), and both Cph1 and Efg1 (HLC54) were spotted onto the indicated S<u>X</u>D media (<u>X</u> = Pro, Arg, Orn or Asp) and incubated at 37 °C for 48 h. **C.** Levels of active GTP bound form of Ras1 (Ras1-GTP) increase upon amino acid induction. Extracts were prepared from pooled WT (PMRCA18) macrocolonies grown for 24 h at 37 °C on the specified S<u>X</u>D medium. The levels of total Ras1 and the activated forms (Ras1-GTP) were determined by immunoprecipitation. **D.** Arginine catabolism is required for arginine-induced morphogenesis. Cells were spotted on S<u>R</u>D (Arg) supplemented with nor-NOHA, a competitive inhibitor of arginase. **E.** Proline catabolism is required for proline-induced morphogenesis. Cells were spotted on S<u>P</u>D (Pro) supplemented with L-THFA, a competitive inhibitor of proline dehydrogenase. For **D** and **E**, macrocolonies (PMRAC18) were grown at 37 °C and photographed after 72 h. Lower images are magnified 2X in comparison to upper images.

Our results regarding the clear Ras1-dependence suggested that amino acid-initiated signals promote GTP-GDP exchange. To test this notion, we assessed the levels of Ras1-GTP in cells after induction by amino acids **(Fig. 2C**). Our results clearly show that in contrast to cells induced with aspartate, cells induced with arginine, proline and ornithine had increased levels of activated Ras1 in its GTP bound form. We attempted to directly assess the requirement of adenylyl cyclase, however, the previously characterized *cdc35*Δ/Δ (*cyr1*) strain [26] did not grow in the synthetic media used here, even when the media was supplemented with 100 μg/ml uridine and/or 10 mM dibutyryl cAMP (data not shown). The lack of growth of this strain, which was contrary to our expectations, precluded a direct assessment of the role of Cyr1.

We tested whether amino acid catabolism was required to activate PKA-signaling by examining the morphology of colonies from cells grown on medium containing arginine or proline as sole nitrogen source and supplemented with enzyme specific inhibitors. In *C. albicans*, arginine is primarily catabolized via the arginase (*CAR1*) pathway commencing with the hydrolysis of arginine to ornithine and urea. N^ω^-hydroxy-nor-arginine (Nor-NOHA), a potent competitive inhibitor of arginase [41], clearly inhibited arginine-induced filamentation in a dose-dependent manner (**Fig. 2D**). Similarly, L-tetrahydrofuroic acid (L-THFA), a specific competitive inhibitor of proline dehydrogenase (Put1)[42, 43], greatly impaired filamentation in a dose-dependent manner (**Fig. 2E**). The data demonstrate that the arginine- and proline-inducing signals are derived from their catabolism.

### Increased ATP resulting from mitochondrial metabolism of morphogenic amino acids activate Ras1/cAMP/PKA signaling

Intracellular levels of ATP are thought to provide a key input for Ras1/cAMP/PKA signaling [14]. Consistent with this notion, in comparison to cells grown in the presence of non-inducing nitrogen sources, such as aspartate and ammonium sulfate, cells grown in the presence of the morphogenic amino acids arginine, ornithine or proline contained similar, and significantly higher levels of ATP (**Fig. 3A**). Urea robustly induces filamentous growth, however, urea-derived signals bypass Ras1; morphogenic induction is dependent on *DUR1,2*-dependent metabolism that generates CO_2_ [30]. Interestingly, cells grown on media with urea, contained significantly lower levels of ATP.

**Fig 3.**
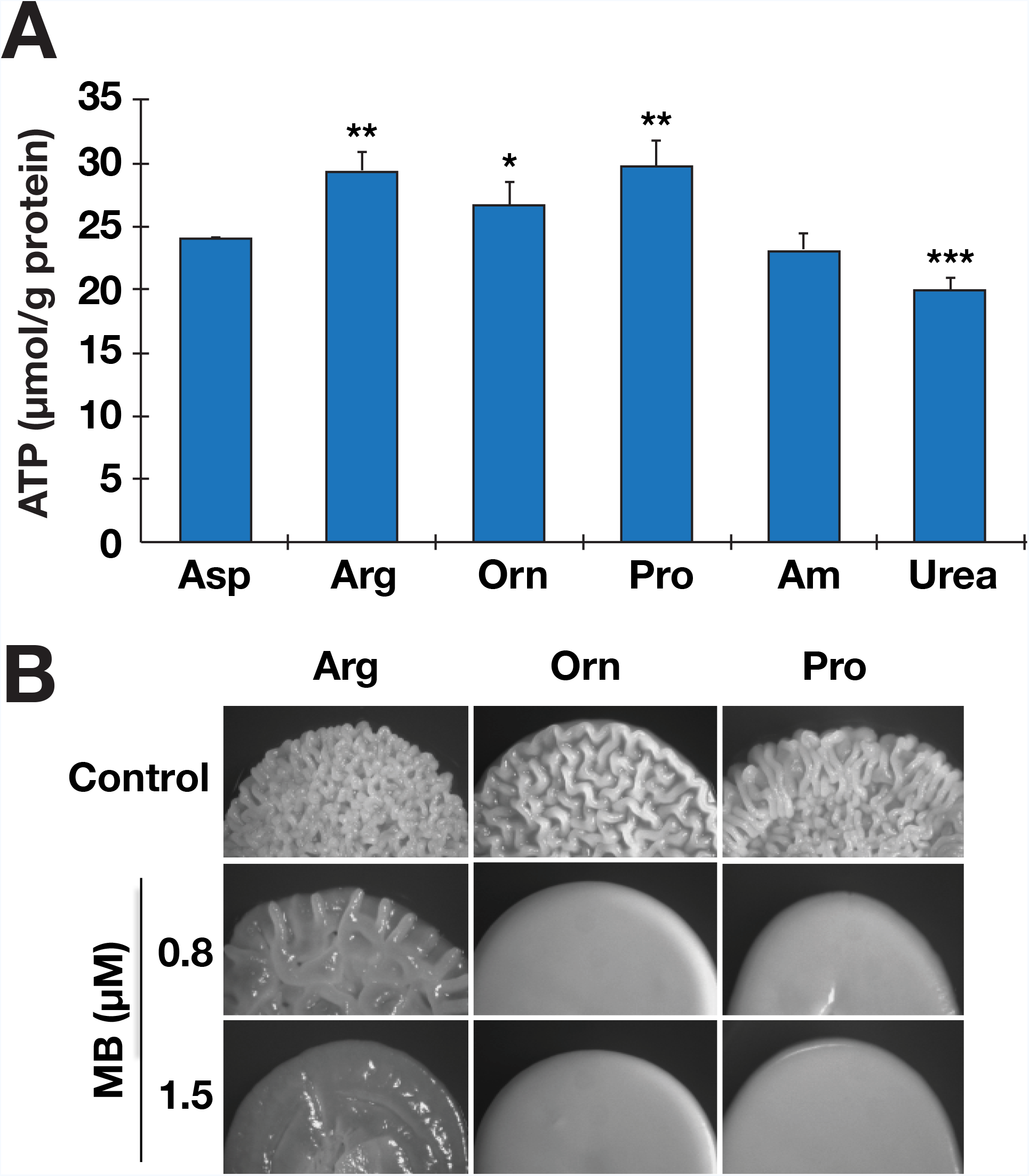
Amino acid-induced morphogenesis requires mitochondrial oxidative phosphorylation. **A.** ATP levels in macrocolonies (PMRCA18) formed 24 h after spotting cells on the indicated S<u>X</u>D medium (X = Asp, Arg, Orn, Pro, Am (ammonium sulfate) or Urea) incubated at 37 °C. The levels of ATP in three biological replicates normalized to total protein are plotted. The values from each biological replicate is the average of 2-3 technical replicates. Statistically significant changes in ATP levels, as compared to cells grown on Asp, are indicated (ave. ± CI; **, p value < 0.01; *, p value < 0.05). **B.** Uncoupling of mitochondria reduces amino acid-induced filamentation. Cells (PMRCA18) were spotted on S<u>X</u>D media (X = Arg, Orn or Pro) supplemented with indicated amount of methylene blue (MB); macrocolonies were grown at 37 °C and photographed after 24 h.

Arginine and ornithine are catabolized to proline in the cytoplasm, and proline is subsequently metabolized to glutamate and then α-ketoglutarate in the mitochondria [12]. These metabolic events generate the reduced electron donors, FADH_2_ and NADH, which are oxidized by the mitochondrial electron transfer chain leading to ATP synthesis (**Fig. 3A**). We posited that the increased levels of ATP resulting from the catabolism of arginine, ornithine and proline is the consequence of their shared metabolic pathway. To test this, we used methylene blue (MB), which uncouples electron transport from the generation of a proton motive force across the inner mitochondrial membrane. The inclusion of MB in media containing ornithine or proline completely inhibited filamentous growth (**Fig. 3B**). Interestingly, the inhibitory effect of MB in cells growing on arginine was not complete, and a higher concentration of MB was required to noticeably inhibit filamentation. These latter findings suggest that an alternative arginine-induced pathway that is independent of ATP-generating mitochondrial metabolism exists in *C. albicans*.

We sought independent means to assess levels of reduced electron donors generated by the metabolism of the morphogenic amino acids. The membrane-permeable redox indicator TTC (2,3,5 triphenyltetrazolium chloride, colorless) is converted to TTF (1,3,5-triphenylformazan, red) in the presence of NADH and has been be used to monitor mitochondrial respiratory activity of colonies [15, 44]. Colonies growing on proline, arginine, and ornithine exhibited a more intense, deep red pigment than colonies growing on aspartate (**Fig. S4, top panel**). The redox-sensitive dye resazurin can be used in liquid culture to monitor the reducing capacity of the intracellular environment [45]; resazurin is non-fluorescent, but is readily reduced by NADH or to a lesser extent by NADPH to highly red fluorescent resorufin (excitation 560 nm, emission 590 nm). Consistent with the results obtained using TTC, cells growing with proline, arginine, or ornithine as sole nitrogen source exhibited 6 – 8-fold more resorufin fluorescence than aspartate grown cells (**Fig. S4, bottom panel**). These results indicate that cells grown in the presence of proline as the sole nitrogen source have a reducing intracellular environment, a finding aligned with the previous report by Land et al. [11].

### Proline metabolism generates the primary signal for arginine-induced morphogenesis

Arginine is degraded in a pathway that bifurcates after the initial reaction catalyzed by Car1, which forms ornithine and urea (**Fig. 4A**). Ornithine is subsequently metabolized by ornithine aminotransferase (*CAR2*) to form glutamate γ-semialdehyde, which spontaneously converts to Δ^1^-pyrroline-5-carboxylate (P5C). P5C is converted to proline by the *PRO3* gene product. Cytoplasmic proline is transported into the mitochondria where it is converted back to P5C by proline oxidase (*PUT1*). Finally, the mitochondrial P5C is converted to glutamate by the *PUT2* gene product (Marczak and Brandriss, 1989; Siddiqui and Brandriss, 1989), which is then converted to α-ketoglutarate via Gdh2. Urea is further catabolized in the cytosol by urea amidolyase (*DUR1,2*) forming NH_3_ and CO_2_.

**Fig 4.**
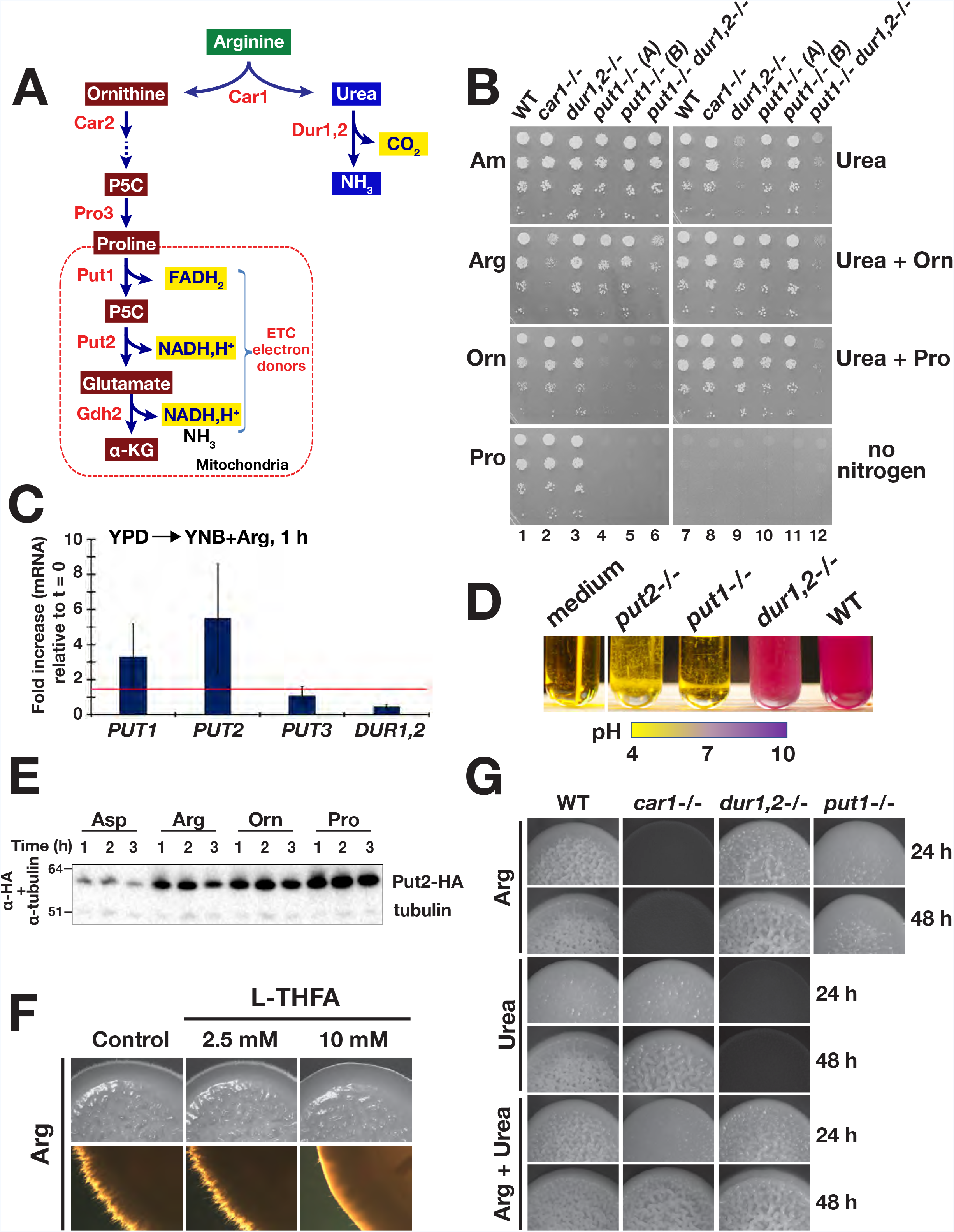
A bifurcated pathway for arginine-induced morphogenesis. **A.** Scheme of arginine catabolic pathway. **B.** Growth-based assays. Five microliters (5 μl) of serially diluted cells were spotted onto the surface of S<u>X</u>D (<u>X</u> = Am (Ammonium sulfate), Arg, Orn, Pro or Urea) and then grown for 48 h at 30°C. Strains used: wildtype (WT; PMRCA18), *car1*-/- (CFG077), *dur1,2*-/- (CFG091), *put1*-/- (A) (CFG122), *put1*-/- (B) (CFG155), and *put1*-/- *dur1,2*-/- (CFG158). **C.** Arginine rapidly derepresses proline catabolic genes. *PUT1*, *PUT2*, *PUT3* and *DUR1,2* expression in wildtype (SC5314) cells 1 h at 37 °C after shifting from YPD (t = 0) to YNB+Arg (pH = 6.0). Gene expression was determined by qRT-PCR using the levels of *RIP1* to normalize expression [96]. **D.** Proline catabolic pathway is required for growth in arginine as a sole nitrogen and carbon source. Cells with the indicated genotypes were harvested from log phase YPD cultures and diluted in YNB+Arg+BCP medium (pH = 4.0) to OD_600_ ≈ 0.01, cultures were incubated for 16 h at 37 °C. Alkalinization (shift to purple) correlates with growth. Strains used: WT (SC5314), *dur1,2*-/- (CFG246), *put1*-/- (CFG149), and *put2*-/- (CFG143). Identical results were obtained using PMRCA18-derived mutants. **E.** Rapid activation of proline catabolism in the presence of arginine, ornithine, and proline. Immunoblot analysis of Put2-HA in whole cell lysates prepared from CFG185 (*PUT2/PUT2-HA*) cells grown at 37 °C in the specified S<u>X</u>D media for the indicated times. Cells were pre-grown in YPD and inoculated at an OD_600_ of 0.5. Levels of α-tubulin were used to control loading. **F.** Pharmacological inhibition of proline dehydrogenase (Put1) reduced filamentous growth of *C. albicans* in the presence of arginine. Macrocolonies of wildtype (WT; PMRCA18) grown at 37 °C for 72 h on S<u>R</u>D medium supplemented with L-THFA as indicated. **G.** Filamentous growth of strains on S<u>X</u>D media (X = Arg, Urea, or Arg + Urea as sole nitrogen sources). Macrocolonies were grown at 37 °C and photographed at 24 and 48 h. Strains used: wildtype (WT; PMRCA18), *car1*-/- (CFG077), *dur1,2*-/- (CFG091), and *put1*-/- (CFG122).

Based on our results demonstrating that the filament-inducing effect of ornithine and proline requires mitochondrial respiration, we investigated if both branches of the bifurcated arginine degradative pathway could independently trigger filamentous growth. To accomplish this, we used a CRISPR/Cas9 strategy to construct PMRCA18-derived strains individually lacking *CAR1* (**Fig. S3D**), *DUR1,2* (**Fig. S3E**), or *PUT1*, *PUT2* and *PUT3* (**Fig. S3F**), or both *PUT1* and *DUR1,2* (**Fig. S3G**). Growth-based assays, on solid and in liquid media, confirmed that the *car1*-/- strain exhibited impaired growth on synthetic glucose medium (S<u>X</u>D) containing arginine as a sole nitrogen source, whereas the strain grew like wildtype (WT) on S<u>X</u>D medium containing either 10 mM ornithine, proline or urea (**Fig. 4B and Fig. S5**). As expected, and similar to previous reports [30], the *dur1,2*-/- strain exhibited severely impaired growth on medium containing urea as sole nitrogen source, but grew well in media containing arginine, ornithine or proline as sole nitrogen sources (**Fig. 4B and Fig. S5**). Cells lacking the proline oxidase (*put1*-/-) were able to grow in media containing arginine, but unable to grow when ornithine or proline were the sole source of nitrogen (**Fig. 4B and Fig. S5**), indicating that ornithine utilization is strictly dependent on the mitochondrial proline catabolic pathway (**Fig. 4B and Fig. S5**).

Interestingly, and quite surprisingly, the *put1*-/- *dur1,2*-/- double mutant strain retained the ability to grow with arginine as sole nitrogen source, albeit slower, clearly suggesting that an arginase-independent arginine utilization pathway exists in *C. albicans* (**Fig. 4B and Fig. S5**). In subsequent growth-based assays the *car1*-/- strain exhibited glucose-dependent growth phenotypes. The *car1*-/- strain did not grow when arginine was present as the sole nitrogen and carbon source, and in the absence of glucose, the *car1*-/- strain did not alkalinize the media (**Fig. S6**). Thus, in the presence of high glucose, the arginase-independent pathway merely enables the use of arginine as a nitrogen source.

Next we analyzed the expression of genes involved in arginine catabolism in cells after shifting them to minimal medium containing 10 mM arginine (YNB+Arg) as sole nitrogen and carbon source (**Fig. 4C**). One hour after the shift, the proline catabolic genes *PUT1* and *PUT2* were significantly upregulated. The levels of *PUT3*, the proline activated transcription factor that is constitutively bound to the promoter of *PUT1* and *PUT2*, did not change [46]. Strikingly, *DUR1,2* gene expression remained constant. Contrary to the assumption that Dur1,2 is responsible for alkalization of the medium, the consequence of the deamination of arginine-derived urea [47], we observed that the *dur1,2*-/- mutant still alkalinized the medium (**Fig. 4D**). Notably, both *put1*-/- and *put2*-/- strains failed to grow in this medium (**Fig. 4D**), indicating that the proline catabolic pathway branch of arginine utilization is essential for growth when arginine is both carbon and nitrogen source. Accordingly, an increased flux through the proline branch of the pathway and subsequent deamination of glutamate provides the likely explanation for the alkalization of the medium (**Fig. 4A**).

Consistent with their ability to support growth, arginine, ornithine and proline induced the expression of HA epitope-tagged Put2 (Put2-HA) (**Fig. 4E**). The induction was rapid, 1 h following the shift from YPD to S<u>X</u>D (X = 10 mM Asp, Arg, Orn or Pro), Put2 expression was derepressed in the presence of arginine and ornithine, almost to the levels observed by the addition of proline. Aspartate did not induce Put2 expression. Together these results indicate that arginine and ornithine are efficiently metabolized to proline, and metabolism associated with proline branch is required for the use of these amino acids as energy sources for growth.

To test whether proline catabolism is required for arginine-induced morphogenesis, we tested first whether morphogenesis in the presence of arginine can be reduced by Put1 inhibitor, L-THFA (Zhu et al., 2002; Zhang et al., 2015). As expected, pharmacological inhibition of Put1 by L-THFA inhibited arginine-induced morphogenesis (**Fig. 4F**). We then carried out a genetic analysis to dissect the pathway triggering filamentous growth in the presence of arginine. Consistent with the existing model for arginine-induced morphogenesis [30], the *car1*-/- strain formed extensively wrinkled colonies in the presence of 10 mM urea comprised mainly of filamentous cells (**Fig. 4G**). However, in comparison to wildtype colonies growing on arginine media, wrinkling was delayed and was first noticeable after 48 h of incubation. On media with an equimolar amount of arginine and urea (Arg + Urea) the *car1*-/- strain developed wrinkled colonies clearly visible after only 24 h. These findings suggest that arginine metabolism via the proline branch induces filamentation more rapidly than the CO_2_ (HCO_3_^-^) generated by the Dur1,2-dependent degradation of urea. Consistent with this notion, colonies formed by the *put1*-/- mutant remained relatively smooth even after 48 h of growth (**Fig. 4G**). In summary, our results indicate that the metabolism associated with proline branch of the arginine degradation pathway generates the primary and most rapid signal of arginine-induced morphogenesis.

### Proline utilization is sensitive to carbon source availability and independent of NCR control

The capacity of proline to stimulate filamentous growth is significantly affected by glucose availability (**Fig. 5A**). In comparison to colonies formed on synthetic media with 10 mM proline containing 2% glucose (SP<u>D</u>), colonies on media containing 0.2% glucose (SP<u>D_0.2%_</u>) exhibited larger feathery zones of hyphal cells emanating around their periphery. These findings are reminiscent of reports that *C. albicans* cells grown on media with methionine as nitrogen source and low glucose exhibit robust filamentation [31]. We considered the possibility that glucose repression of mitochondrial function, known to occur in *Saccharomyces cerevisiae* [48, 49], may underlie the difference. Consistent with this notion, macrocolonies formed on SP<u>D</u> were deeper red in color when overlaid with TTC than macrocolonies formed on SP<u>D_0.2%_</u> or on media with 1% glycerol (SP<u>G</u>). The lighter red color of macrocolonies on low glucose, or on glycerol, confirm that cells have lower intracellular levels of NADH, i.e., under derepressing conditions when mitochondria can efficiently oxidize NADH (**Fig. 5A**).

**Fig 5.**
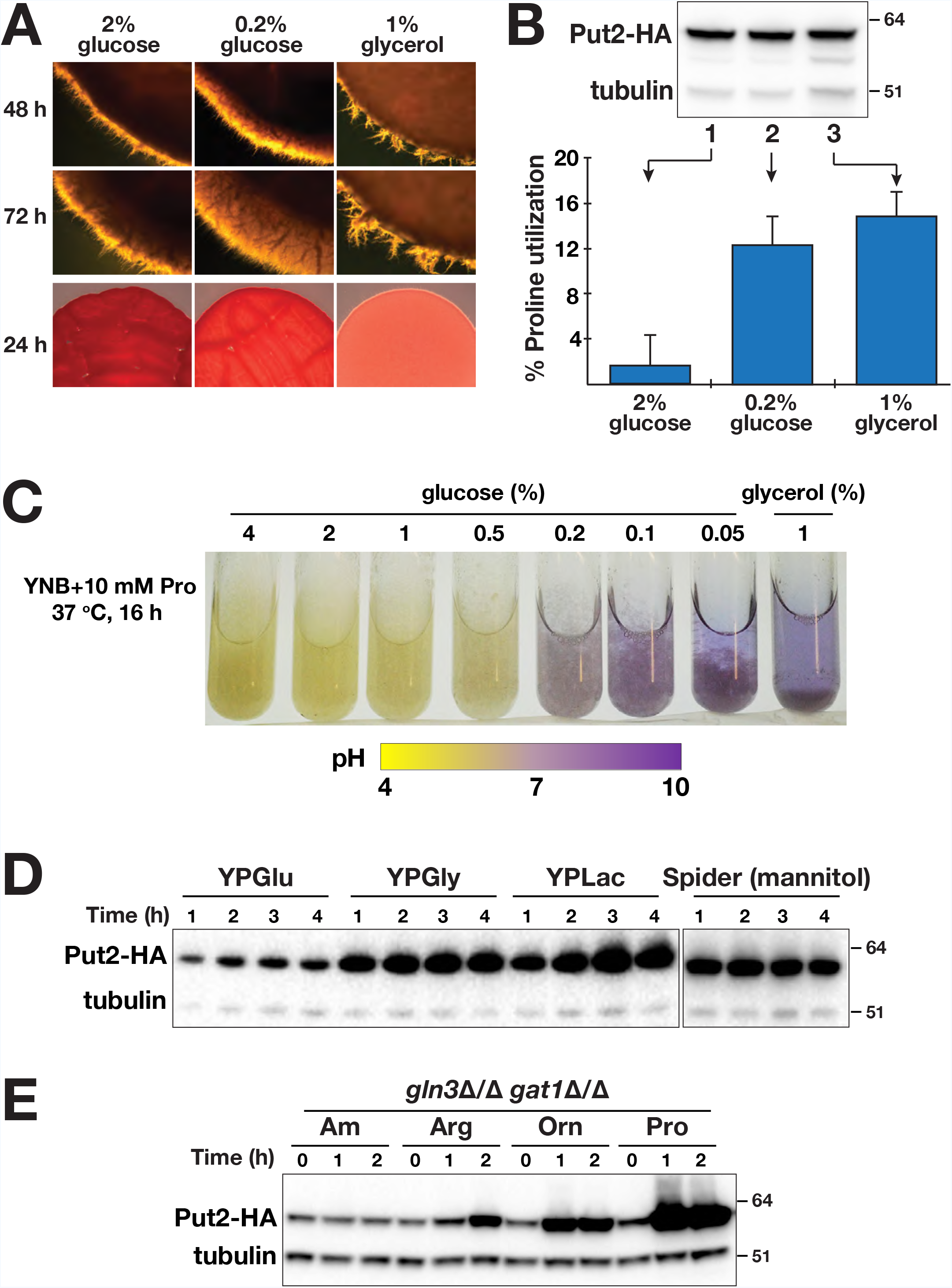
Proline utilization is influenced by carbon source but not by NCR. **A.** Filamentous growth is more robust at lower glucose level. Wildtype cells (PMRCA18) from overnight YPD liquid cultures were washed and then adjusted to OD_600_ of 8, 10 μl aliquots were spotted on media containing 10 mM of proline and the indicated levels of carbon source. Plates were incubated at 37 °C and photographed at 24, 48 and 72 h. TTC overlay assay was performed on a replica plate containing 24 h-old macrocolonies (shown in 24 h panel). **B.** Proline utilization is higher under glucose limited conditions. The levels of proline remaining in culture supernatants of WT (PMRCA18) after 2 h of growth at 37 °C in the presence of different carbon source, as indicated. Results shown are from 5 biological replicates (Ave. ± CI, 95% CL; ***, p value < 0.001). Immunoblot analysis of Put2-HA and α-tubulin (loading control) in cell extracts prepared from CFG185 (*PUT2/PUT2-HA*) grown under identical conditions (inset). **C.** Respiratory growth predominates as glucose level decreases. WT cells were diluted to OD_600_ of 0.5 in pre-warmed synthetic proline media containing 10 mM proline (YNB+Pro+BCP) and the indicated levels of glucose with the initial pH adjusted to 6.0. Cultures were grown for 16 h under vigorous agitation at 37 °C prior to photographing the culture tubes. Glycerol was used as respiratory growth control. **D.** Put2 is highly expressed in cells grown in the presence of non-glucose carbon sources. Immunoblot analysis of cell extracts prepared from CFG185 (*PUT2/PUT2-HA*) cells grown in YP<u>Glu</u> (YP+2% glucose = YPD), YP<u>Gly</u> (YP + 1% glycerol), YP<u>Lac</u> (YP + 1% lactate), or Spider medium (with 1% mannitol) at 37 °C for the timepoints indicated. Cells were pre-grown in YPD and inoculated at an OD_600_ of 0.5. **E.** Proline utilization is insensitive to NCR. Immunoblot analysis of cell extracts prepared from CFG184 (*gln3*Δ*/*Δ *gat1*Δ*/*Δ *PUT2/PUT2-HA*) grown at 30 °C in SD_0.2%_ medium, which contains 10 mM ammonium sulfate (Am) and 0.2% glucose, supplemented with 10 mM of the nitrogen sources and harvested at the timepoints as indicated. Cells were pre-grown in SD and then subcultured to log phase in SD_0.2%_.

We tested the notion that at low glucose concentrations, i.e., non-repressing conditions, cells use proline as a carbon source. Proline utilization was assayed directly by measuring the amount of residual proline in culture supernatants after a 2 h incubation period. In media containing 2% glucose, cells took up < 2% of the proline. By contrast, cells growing in low glucose (0.2%) or 1% glycerol used 12 – 15% of the available proline (**Fig. 5B**). The expression of Put2 was independent of glucose, as the levels of Put2 were similar (**Fig. 5B, insert**). These results indicate that proline is taken up and metabolized more efficiently in cells under non-repressing conditions, suggesting that mitochondrial activity is subject to glucose repression.

To critically test this, we assessed the effect of varying the glucose concentration from 0.05 - 4%. Cells were grown for 16 h in media containing the pH indicator bromcresol purple. At high glucose concentrations (0.5 – 4%) the media remained acidic, indicating cells were growing fermentatively using proline merely as a nitrogen source (**Fig. 5C**). By contrast, at glucose concentrations ≤ 0.2%, the media became alkaline, indicating that cells were respiring and using proline as the primary energy source. The increased flux through the proline pathway is expected to yield elevated NH_3_ generated by the mitochondrial glutamate dehydrogenase (*GDH2*) catalyzed deamination of glutamate. To directly assess mitochondrial activity under these conditions, we carried out extracellular oxygen consumption analysis in a high-throughput microplate format (**Fig. S7**). Cells grown in repressing SP<u>D</u> had the lowest oxygen consumption whereas those grown at SP<u>D_0.2%_</u> had the highest, higher than cells grown in SP<u>G</u>. As previously pointed out, Put2 levels were similar across all conditions (**Fig. 5B**). Together, these results indicate that proline is taken up and then metabolized more efficiently in cells growing under low glucose concentrations. Consistent with this finding, Put2 expression was derepressed in rich media containing yeast extract and peptone when non-repressing, non-fermentative carbon sources replaced glucose; i.e., glycerol or lactate (**Fig. 5D**). Similarly, cells express elevated levels of Put2 when grown in hyphal inducing Spider medium, a medium rich in amino acids and mannitol as a primary carbon source.

Nitrogen regulation of transcription in fungi is a suprapathway response that is commonly referred to as nitrogen catabolite repression (NCR), which functions to ensure that cells selectively use preferred nitrogen sources when available. Briefly, NCR regulates the activity of GATA transcription factors Gln3 and Gat1; in the presence of preferred nitrogen sources, these factors do not gain access to the promoters of NCR-regulated genes (reviewed in[50]). Previous studies have shown that certain amino acids, traditionally classified as poor (e.g., proline) in *S. cerevisiae*, were readily utilized by *C. albicans* mutants lacking Gln3 and Gat1 [51]; the introduction of null alleles of both *GLN3* and *GAT1* in *C. albicans* did not impair growth using proline as sole nitrogen source, whereas growth on urea was severely affected. Consistent with these findings, we found that Put2-HA was constitutively expressed in *gln3*Δ/Δ/ *gat1*Δ/Δ mutant grown in medium containing high levels of the preferred nitrogen source ammonium sulfate (**Fig. 5E**). Our data indicate that in *C. albicans* proline utilization is not subject to NCR, a conclusion aligned with recently published transcriptome analyses[46].

### Proline induces hyphal growth within phagosomes and enables *C. albicans* to escape from engulfing macrophages

We sought to place our novel insights regarding the critical role of proline metabolism in the induction of hyphal growth in a broader biological context and tested whether proline catabolism affects the capacity of *C. albicans* cells to form hyphae within macrophages and escape killing. First, using indirect immunofluorescence microscopy we examined whether Put2-HA is expressed in *C. albicans* cells engulfed by murine RAW264.7 macrophages (**Fig. 6A**). *C. albicans* CFG185 (*PUT2/PUT2-HA*) cells were co-cultured with macrophages (MOI of 5:1; C:M) for 90 min. Strain CFG185 exhibits activation of proline catabolism in the presence of arginine, ornithine, and proline (**Fig. 4E**). The macrophages were imaged using antibodies against the HA tag (1°, rat anti-HA; 2°, goat anti-rat antibody conjugated to Alexa Fluor 555) and LAMP-1, a lysosomal marker that is enriched in phagosomes. Confocal images clearly showed that *C. albicans* cells engulfed by macrophages express Put2, and that the Put2 expressing fungal cells localized to Lamp1 compartments (see the orthogonal view of merged channels, lower left panel). The results indicate that *C. albicans* cells within macrophage phagosomes express Put2.

**Fig 6.**
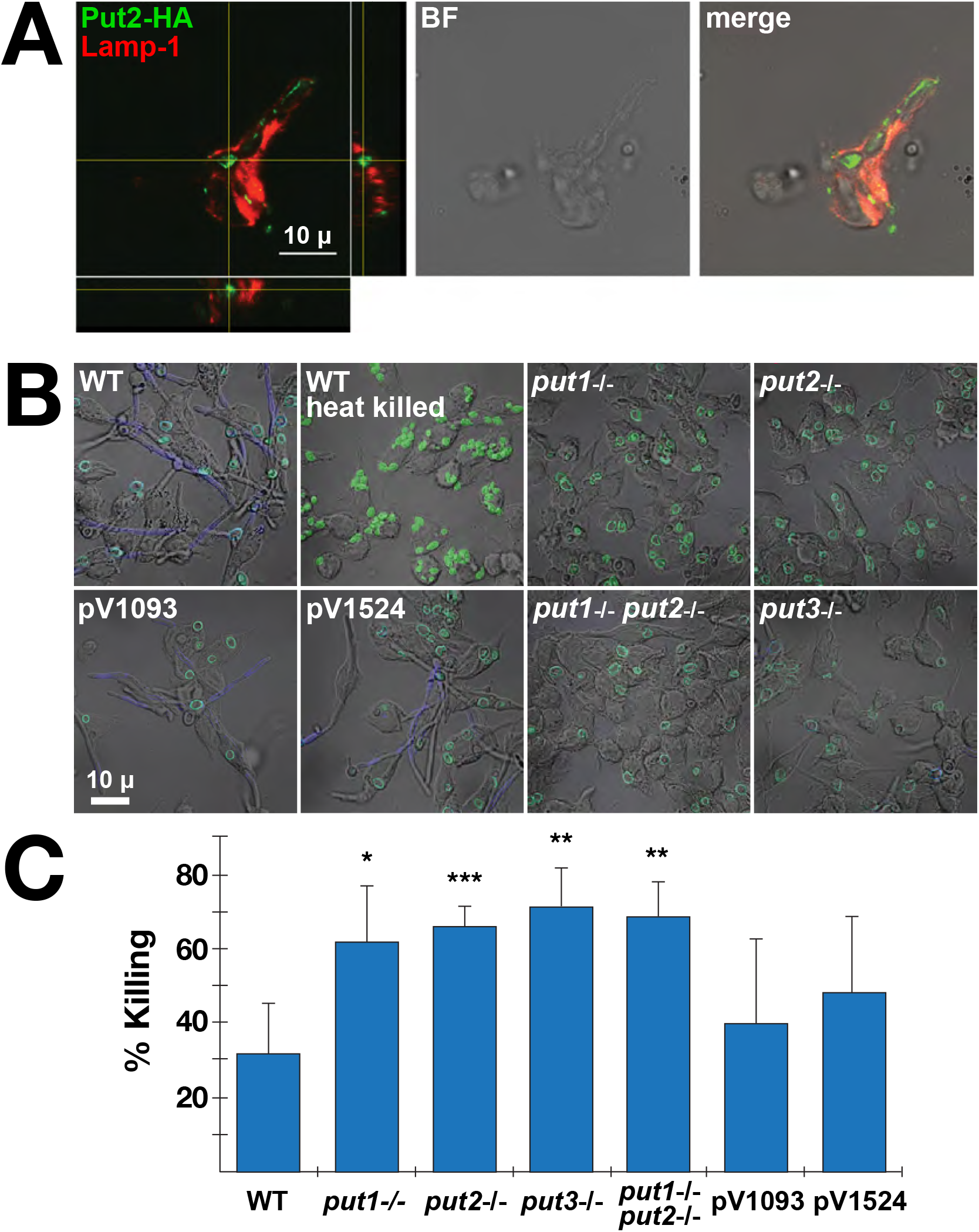
Mitochondrial proline catabolism is required by *C. albicans* cells to escape murine macrophages. **A.** Confocal immunofluorescence microscopy of *C. albicans* cells expressing Put2-HA (CFG185) in the phagosomes of RAW264.7 macrophages. Primary antibodies (rat anti-HA and rabbit anti-Lamp1) and secondary antibodies (Alexa Fluor 555 conjugated goat anti-rat antibody and Alexa Fluor 488 conjugated goat anti-rabbit) were used to visualize Put2 and the lysosomal compartment, respectively. Orthogonal view of merged channels is shown in the lower right panel. Scale bar = 10 μ. **B.** Proline catabolism is required for hyphal growth of *C. albicans* in macrophages. Wildtype (WT; SC5314), heat killed WT, *put1*-/- (CFG139), *put2*-/- (CFG207), *put3*-/- (CFG146), *put1*-/- *put2-/-* (CFG159) and CRISPR/Cas9 control strains CFG181 (pV1093) and CFG182 (pV1524) pre-grown in YPD and stained with FITC were co-cultured with RAW264.7 macrophages at a MOI of 3:1 (C:M). After 30 min, external non-phagocytosed cells were removed by washing, and the co-cultures were incubated an additional 4 h. External (escaping) hyphal cells were stained with calcofluor white (CFW). Scale bar = 10 μ. **C.** Macrophage killing of *C. albican*s. Strains as in **B** were co-cultured with RAW264.7 at a MOI of 3:1 (C:M) for 3 h. After lysing macrophages, viability of *C. albicans* was assessed by quantitating the number of CFU. The percent killing was determined by comparison to the viability of cells grown in the absence of macrophages.

Next, we assessed the importance of the proline catabolic pathway components in the capacity to escape macrophage. To facilitate comparisons with results obtained in other laboratories, we repeated the construction of the proline catabolic pathway mutations in the SC5314 strain background; strains lacking *PUT1*, *PUT2*, *PUT3* or both *PUT1* and *PUT2* were constructed using CRISPR/Cas9. The full genome of each mutant strain was sequenced; the sequence coverage varied from 42 – 65X and after assembly the contig coverage accounted for ≥98 of the reference SC5314 genome (Assembly 22, version s06-m01-r01; [52]). Each strain was found to carry the intended null mutation in the correct chromosomal locus and no large scale dissimilarities to the reference genome or off-target mutations were evident. Furthermore, no phenotypic differences were detected in comparison to the PMRCA18-derived strains (data not shown).

As expected, SC5314 (WT) and CRISPR/Cas9 control strains (pV1093 and pV1524), lacking guide sequences to target Cas9, exhibited robust hyphal growth when co-cultured with RAW264.7 macrophages (**Fig. 6B**). By contrast, and similar to heat killed SC5314, the strains carrying *put1*-/-, *put2*-/-, *put3*-/- and *put1*-/- *put2-/-* mutations were unable to efficiently form filaments from within engulfing macrophages (**Fig. 6B**). As hyphal formation enables *C. albicans* cells to escape macrophages and thereby facilitates survival, we analyzed the candidacidal activity of macrophages by assessing fungal cell viability by assessing colony forming units (CFU). Consistent with our microscopic analysis, in comparison to wildtype cells, the proline mutants were killed more efficiently (**Fig. 6C**). Together, these results indicate that *C. albicans* cells rely on proline catabolism to induce hyphal growth in phagosomes, a response that facilitates escape from killing by macrophages.

## Discussion

In this study we have found that ATP generating mitochondrial proline catabolism is required to induce hyphal development of *C. albicans* cells in phagosomes of engulfing macrophages. The finding that proline catabolism, also required for the utilization of arginine and ornithine, is required to sustain the energy demands of hyphal growth provides the basis to understand the central role of mitochondria in fungal virulence. The energy status of the fungal cell is clearly a key signal that engages the genetic programs underlying yeast-to-hyphal transitions. The dependence on the energy producing proline catabolic pathway to induce *C. albicans* cells to switch morphologies is instrumental in their ability to escape from macrophages. Our results are consistent with a recent model postulating that elevated cellular levels of ATP induces hyphal morphogenesis [14] and with early reports that amino acid catabolism promotes filamentous growth [12, 13, 53]. Our experimental findings are schematically summarized in Fig. 7.

**Fig 7.**
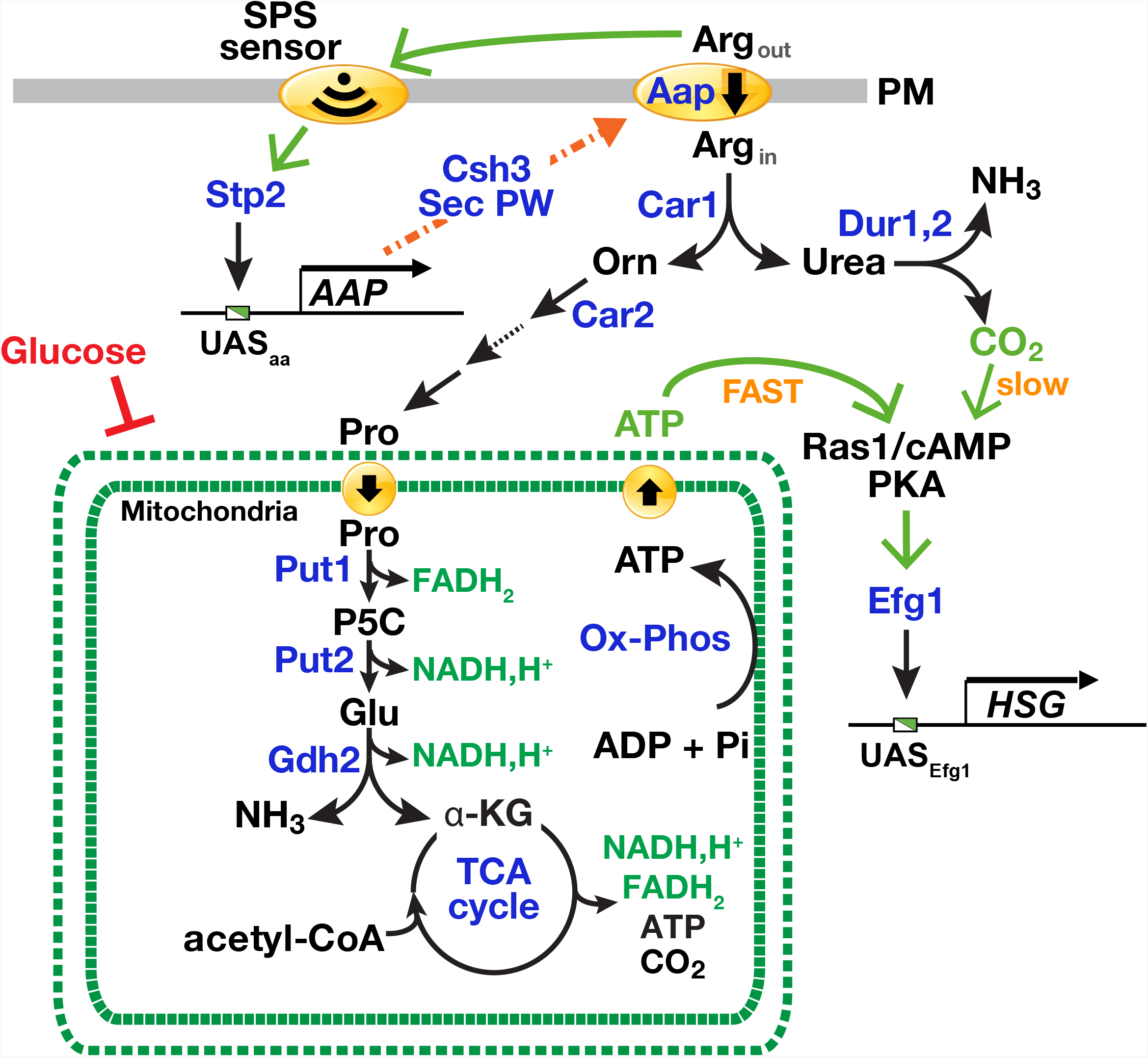
Arginine induces morphogenesis in *C. albicans* through mitochondria-dependent activation of Ras1/cAMP/PKA pathway. The presence of extracellular arginine enhances arginine uptake by binding to the SPS-sensor, leading to the endoproteolytic activation of transcription factor Stp2. The active form of Stp2 efficiently targets to the nucleus and binds the UAS_aa_ in the promoter of genes encoding amino acid permeases (*AAP*). Amino acid permeases are cotranslationally inserted into the membrane of the ER, and transported to the plasma membrane (PM, arrow) via the secretory pathway, a process that requires the ER membrane-localized chaperone Csh3. The increased functional expression of amino acid permeases (Aap) lead to an enhanced capacity to take up arginine. Intracellular arginine is catabolized by arginase (Car1) yielding ornithine and urea. Urea is further degraded by the urea amidolyase (Dur1,2) generating CO_2_ and NH_3_. Ornithine is further catabolized to proline in the cytoplasm in a series of enzymatic reactions starting with the ornithine aminotransferase (Car2). Proline is transported into the mitochondria where it is catabolized by Put1 and Put2 forming glutamate. These reactions generate the reduced electron carriers FADH_2_ and NADH,H^+^, respectively. Glutamate is oxidized by glutamate dehydrogenase (Gdh2) forming α-ketoglutarate in a reaction that liberates NH_3_ and generates NADH,H^+^. α-ketoglutarate feeds directly into the mitochondria-localized TCA cycle. The reduced electron carriers generated by proline, glutamate and TCA cycle metabolic events are oxidized in reactions coupled to the generation ATP by mitochondrial oxidative phosphorylation. The elevated levels of ATP in the cytoplasm activate the adenyl cyclase (Cyr1) in a Ras1-dependent manner, which activates the downstream protein kinase A (PKA) signaling pathway and the effector transcription factor Efg1. The active phosphorylated form of Efg1 binds the UAS_*Efg1*_ in the promoter of hyphal specific genes (*HSG*), thereby inducing yeast-to-hyphal morphogenesis. The catabolism of arginine via the proline pathway induces hyphal growth more rapidly (FAST) than the Dur1,2 generated CO_2_ (slow). Mitochondrial activity is repressed in the presence of high glucose.

Our work provides a framework to integrate several fragmentary observations regarding amino acid-induced morphogenesis. For example, Land et al. [11, 12] observed that the most potent morphogenic amino acids arginine and proline are those metabolized to glutamate. Our results show that this occurs strictly via the mitochondrial localized proline utilization pathway essentially as described in *S. cerevisiae* [54–56] with the exception that proline metabolism is not under nitrogen regulation (**Fig. 5E**,[46]. Consistently, ornithine, an intermediate in arginine catabolism, also acts as a potent inducer of morphogenesis (**Fig. 1A**; [12, 53]. Glutamate is further converted to α-ketoglutarate, an intermediate in the TCA cycle. These metabolic reactions are coupled to the generation of reduced electron carriers FADH_2_ and NADH, which are oxidized in the mitochondria powering ATP synthesis. Amino acid induction of hyphal growth exhibits a strict requirement for Ras1 (**Fig. 2B**) and cells grown in the presence of these inducing amino acids have high levels of active Ras1 (**Fig. 2C**) and elevated levels of intracellular ATP (**Fig. 3A**). The metabolic inhibitors nor-NOHA (Car1) and L-THFA (Put1) and the mitochondrial uncoupler methylene blue (MB) block the induction of filamentation (**Fig. 2D, E** and **Fig. 3B**). Our analysis demonstrates that arginine and proline induce morphogenesis by virtue of a shared metabolic pathway (**Fig. 4C-F**).

Together, our findings are well aligned to the recent model proposed by Grahl et al. [14], where mitochondrial ATP synthesis facilitates Ras1 activation in cooperation with the adenylate cyclase (Cyr1) leading to increased cAMP production and to activation of the Efg1 transcription factor. The finding that arginine-induced hyphal growth occurs rapidly (**Fig. 4G**), suggests that a brief exposure to arginine may suffice to trigger filamentous growth. According to Grahl et al. (2015), Ras1 activation by ATP appears to be independent of the AMP kinase, a key regulator of cellular energy homeostasis. The ATP-binding pocket within the active site of mammalian adenylyl cyclase has been shown to act as an ATP sensor [57]. Although it has been proposed that Cyr1 may function similarly as an ATP sensor this has yet to be confirmed in *C. albicans*. Regardless of the mechanism, exceeding a critical threshold of ATP is likely required to induce cAMP synthesis. It is known that the cAMP produced by Cyr1 does not necessarily correlate to the strength of the inducer and that transient short-lived spikes in cAMP are sufficient to trigger phosphorylation and eventually activation of Efg1 [27]. Consequently, spikes of ATP transiently generated by proline catabolism may efficiently induce hyphal specific genes (*HSG*).

We have clearly shown that arginine-, ornithine- and proline-induced hyphal growth is dependent on Ras1, which is not accounted for by other models of amino acid-induced morphogenesis (reviewed in [4, 7], despite the fact that Ras1 is known to be important in induction of filamentous growth in the presence of amino acid-rich serum [22]. Both the presumed amino acid sensitive Gpr1-Gpa2 pathway [58, 59] and the Dur1,2-dependent CO_2_ model for arginine-induced morphogenesis [30] are thought to bypass Ras1 and involve direct interactions with adenylyl cyclase (Cyr1). Also, contrary to the previous report [30], CO_2_ generated by the Dur1,2-dependent catabolism of urea is not the primary morphogenic signal. Specifically, induction of filamentous growth in the presence of arginine or proline as sole nitrogen source proceeds more quickly than that observed by the metabolism of urea (**Fig. 4G**). In addition, *DUR1,2* expression is tightly regulated by NCR, i.e., in the presence of ammonia, urea metabolism is repressed [51]. By contrast, the conversion of arginine to proline is not under NCR control (**Fig. 5E**,[46]). Finally, when cells were shifted from YPD to medium containing arginine as sole carbon and nitrogen source, proline catabolic genes (*PUT1* and *PUT2*) were derepressed much faster than *DUR1,2* (**Fig. 4C**), indicating that arginine is rapidly converted to proline. We have noted that the constitutive expression of arginase represents a common and undesired technical problem in proteomic analyses using SILAC (Stable Isotope Labeling by/with Amino acids in Cell culture) due to the rapid conversion of arginine to proline in eukaryotes [60–63]. In *Schizosaccharomyces pombe*, the deletion of two arginase genes (one a *CAR1* homologue) and the single ornithine transaminase (*CAR2* homologue) rectified this problem [60]. We predict, that similar deletions would be helpful in the quantitative analysis of the *C. albicans* proteome.

Earlier reports by Nickerson and Edwards [64] and Land et al. [11] suggested that mitochondrial activity is repressed during filamentous growth. By contrast, other more recent work has shown that hyphal formation occurs predominantly under aerobic conditions [16] and is associated with increased respiratory activity [14, 15]. Based on our findings (**Fig. 5C**), the seemingly conflicting observations could be explained if, as in *S. cerevisiae*, the synthesis of mitochondrial respiratory enzymes are subject to glucose repression [65, 66]. There is surprisingly little information available regarding glucose repression of mitochondrial function in *C. albicans*, and whether the regulatory circuits are wired similar to those in *S. cerevisiae*. However, we note that Land et al. [11] used growth conditions with high glucose (~1.8%; 100 mM), whereas studies by [14, 15] were carried out using low glucose (10 mM, i.e., ≈ 0.2%).

In striking constrast to the current view that *C. albicans* mitochodrial function is insensitive to glucose repression [67–69], our results clearly demonstrate that glucose represses respiration in the presence of proline (**Fig. S7**). Cells grown aerobically in high glucose exhibit fermentative metabolism (**Fig. 5C**), i.e., the well-characterized Crabtree effect [70]. In glycolysis, conversion of glucose to pyruvate is coupled to reduction of NAD^+^ and to the generation of ATP. Only small amounts of the cofactor is available in the cytosol. Consequently, when mitochondrial functions are glucose repressed, cells use fermentation to oxidize NADH and regenerate NAD^+^, thereby enabling cytoplasmic ATP synthesis to continue. Under conditions when proline is the sole nitrogen source and high glucose is present, cells use glucose for energy and as carbon-source, whereas proline catabolism merely supplies cells with nitrogen, i.e., proline utilization is low (**Fig. 5B**). However, when glucose becomes limiting (<0.2%), the respiratory capacity of mitochondria increases (**Fig. S7**), enabling cells to efficiently oxidize NADH and generate ATP by oxidative phosphorylation; under these conditions cells use proline for energy and as the carbon- and nitrogen-source, i.e., proline utilization is high (**Fig. 5B**). Together our results show that proline metabolism is a sensitive indicator of mitochondrial function in *C. albicans*.

Our observation that high glucose represses mitochondrial function, provides a mechanistic understanding of how high glucose inhibits hyphal morphogenesis [13, 31]. Cells grown on 2% glucose have elevated levels of reduced cofactors, such as NADH (**Fig. 5A**), suggesting that the capacity of mitochondria to oxidize NADH is suboptimal, i.e., the cellular capacity to regenerate NAD^+^ is rate limiting, a phenomenon termed over-flow metabolism [48]. It is important to note that, based on the *S. cerevisiae* paradigm, the pyruvate formed in glycolysis needs to be converted to acetyl-CoA to prime the TCA cycle. The mitochondrial-localized pyruvate dehydrogenase complex is predominantly responsible for the conversion of pyruvate to acetyl-CoA during glucose-limited, respiratory growth [65, 66]. Indeed, pharmacological inhibition of glycolysis has been shown to arrest filamentous growth of *C. albicans* even in the presence of proline [11]. Alternatively, β-oxidation of lipids may contribute the necessary acetyl-CoA [9].

We have placed the SPS sensing pathway, the primary sensing system of extracellular amino acids, in context to the major intracellular signaling pathways governing in nutrient regulated morphogenesis. SPS-sensor initiated signals do not directly induce hyphal growth, but rather facilitate morphogenesis by up-regulating the capacity of cells to take up inducing amino acids (**Fig. 7**). Experimental support for this conclusion includes the following observations. First, amino acid-induced activation of SPS-sensor signaling does not strictly correlate with the induction of filamentous growth (**Fig. 1A**). Second, the inability of a *ssy1* null mutant to undergo morphogenesis can be rescued by expressing a constitutively active form of Stp2 (*STP2*\*) but not Stp1 (*STP1*\*). Stp2 is the effector transcription factor that controls amino acid permease gene expression, whereas Stp1 activates the expression of secreted aspartyl proteases and oligopeptide transporters [34]. Consistently, and similar to Kraidlova et al. [71], we found that the expression of six *C. albicans* orthologues (*GAP1*-*GAP6*) of the *S. cerevisiae* general amino acid permease (*GAP1*) are regulated by the SPS sensing system, perhaps with the exception of *GAP4* expression, which is comparatively expressed at very low levels (data not shown). Third, filamentous growth is dependent on amino acid catabolism. The weak filamentation observed in the *csh3*∆/∆ mutant grown in 10 mM proline can be attributed to the residual uptake of proline as previously described [35]; apparently, the residual systems are expressed and function at high extracellular concentrations of proline [53, 72]. Thus, the filamentous growth defect observed in cells lacking a functional SPS sensing pathway, i.e., *SSY1* or *CSH3* null mutants, is due to the inability to efficiently take up inducing amino acids from the extracellular environment, a requisite for their metabolism [35, 36].

Together our findings have important implications on understanding how *C. albicans* cells interact with host immune cells. Transcriptomic studies examining macrophage-*C. albicans* interactions by Lorenz et al. [9]showed that arginine biosynthesis genes are peculiarly upregulated in phagocytosed cells. Furthermore, the results suggest that the phagosome is likely a glucose-poor environment as an increased expression of genes that favor gluconeogenesis and mitochondrial function was also noted [9]. Interestingly, arginine utilization appears to proceed concomitant with arginine biosynthesis as deduced from the increased arginase transcripts in phagocytosed cells [9, 30]. In a follow-up study, the apparent upregulation of arginine biosynthesis was suggested to be a response to the macrophage oxidative burst [73]. Interestingly, the expression of *DUR1,2* in phagocytosed cells was not significantly altered. Our finding that the enzymes responsible for proline utilization are upregulated indicates that proline is either made available by the host or is the result of arginine catabolism.

In the light of these results, the challenging question is where the hyphae inducing amino acids come from, from the macrophage or from nutrients stored within *C. albicans* cells prior to their being phagocytosed. In *S. cerevisiae*, > 90% of free arginine is sequestered in the vacuole and the non-compartmentalized and cytosolic arginine is catabolized by arginase [74]. Given that arginine is catabolized to proline via the arginase pathway with ornithine acting as a transitory intermediate, it is possible that vacuolar stores of arginine are activated in the phagosome to support the demand for cellular energy. When glucose becomes limiting, *C. albicans* may rely on the catabolism of amino acids, particularly proline, as primary energy. This is reminiscent of the requirement of proline catabolism for Trypanosome survival in the Tsetse fly vector [75–78].

Proline-induced morphogenesis is repressed under acidic conditions [13, 53], presumably a condition confronting newly phagocytized *C. albicans* cells. This raises the interesting conundrum as to how *C. albicans* cells deal with this environmental challenge and filament. It is possible that Stp2-mediated alkalization of the phagosome reported by Vylkova and Lorentz [79] is a key predisposing event that facilitates proline-induced morphogenesis. We found that alkalization is not Dur1,2-dependent (**Fig. 4D**), indicating that an alternative mechanism triggers alkalization. Accordingly, the Stp2-dependent induction of arginine uptake and its subsequent Put1- and Put2-dependent metabolism generates glutamate, which is deaminated to α-ketoglutarate by glutamate dehydrogenase (Gdh2) (**Fig. 7**). The resulting NH_3_ may provide the explanation for the observed alkalization. As already pointed out, the source of amino acids in the macrophage phagosome remains a very interesting question. Numerous metabolic signatures appear to reflect a microenvironment with a poor nitrogen content. For example, based on the transcriptional analysis of the *C. albicans*-macrophage interaction, *OPT1*, encoding an oligopeptide transporter, is upregulated in phagocytosed cells [9]. *OPT1* expression is controlled by the SPS-sensor signaling and the downstream transcription factor Stp1 [34, 37]. *STP1* expression is itself under tight NCR control [37]. Thus, the upregulated expression of *OPT1* strongly suggests that NCR is relieved in phagocytosed cells and that sufficient levels of amino acids are present to induce the SPS-sensor. As to the origin of amino acids in the phagosome, *C. albicans* may excrete amino acids liberated from storage compartments, loaded during growth in rich media. In *S. cerevisiae*, under defined conditions, amino acids are known to be excreted at detectable levels [80] and under certain circumstances activate SPS-sensor signaling [81]. Thus, amino acids may provide an autocrine function to induce filamentous growth of phagocytosed *C. albicans* cells.

The results presented here provide a clear example of how *C. albicans* cells sense and respond to nutrients present in the host to ensure proper nutrient uptake and continued survival. The molecular components underlying nutrient uptake are often referred to as virulence factors. When afforded the opportunity, *C. albicans* will alter developmental programs to optimize nutrient uptake systems that enable the better exploit host environments and to evade the primary immune response [3, 82, 83]. The identification and understanding of fungal virulence factors is necessary to therapeutically disturb their function upon infectious growth and thereby facilitate the ability of host immune systems to re-establish and maintain the integrity of the host. We are excited by the prospect of exploiting mitochondrial proline metabolism to probe the nutrient environment of the macrophage phagosome, a currently poorly characterized environment.

## Materials and methods

### Strains, media and chemicals

*C. albicans* strains are listed in Supplementary Information Table S1 and all primers used are listed in Table S2. All strains were cultivated in YPD medium (1% yeast extract, 2% peptone, 2% glucose) at 30 °C. Minimal synthetic dextrose (SD) medium containing 0.17% YNB (Yeast Nitrogen Base without amino acids and without ammonium sulfate; Difco^TM^), 2% glucose, and 5 g/l ammonium sulfate (≈ 38 mM) was used as indicated. Media were made solid by 2% (w/v) Bacto agar. Where appropriate, 100 or 200 μg/ml nourseothricin (Nou; Jena Biosciences, Jena, Germany) was added to the medium. The ability of amino acids to induce filamentous growth was determined on buffered solid synthetic (S<u>X</u>D) media containing 0.17% YNB, 2% glucose, and 10 mM of the indicated amino acid (<u>X</u>) as sole nitrogen source, or at concentrations as described in the figure legends. Fifty mM 2-(N-morpholino) ethanesulfonic acid (MES) was included in media and the pH was adjusted to 6.0 using NaOH. To minimize residual nitrogen, the S<u>X</u>D media were made solid using 2% (w/v) highly purified agar (Biolife, Milano, Italy). Where indicated 0.2% glucose, 1% lactate or 1% glycerol replaced 2% glucose as carbon source. The following media were used to screen CRISPR/Cas9-derived knockout phenotypes: YPD-MM; S<u>U</u>D; S<u>P</u>D; and YNB+Arg+BCP. YPD-MM is standard YPD supplemented with 1.5 mg/ml MM (2-([(([(4-methoxy-6-methyl)-1,3,5-triazin-2-yl]-amino)carbonyl)amino]-sulfonyl)-benzoic acid; Dupont™ Ally); S<u>U</u>D and S<u>P</u>D were prepared as S<u>X</u>D containing urea (<u>U</u>), or proline (<u>P</u>) as sole nitrogen source; YNB+Arg+BCP contains 0.17% YNB, 10 mM arginine (Arg) as sole nitrogen and carbon source, and 0.03 μg/mL bromocresol purple (BCP; Sigma) as indicator, with the pH adjusted to 4.0 using 1 M HCl. Growth in the presence of specific metabolic inhibitors was assessed on media containing nor-NOHA (N-hydroxy-nor-L-arginine; BioNordika AB, Sweden) prepared in 100% dimethyl sulfoxide (DMSO) as 56 mM concentrated stock; a 26 mM working stock was prepared freshly diluting in ddH_2_O. L-tetrahydro-2-furoic acid (L-THFA; Sigma) and methylene blue (MB; Sigma), were freshly prepared in ddH_2_O as 1 M and 3 mM stocks, respectively. *Escherichia coli* strain DH10B™ was used for the construction of plasmids; LB medium supplemented where required with carbenicillin (Cb, 50 μg/ml), Nou (50 μg/ml), and/or chloramphenicol (Cm, 30 μg/ml). LB was made solid by 1.5% Bacto agar. Liquid cultures were grown with agitation at 150-200 rpm. The density of yeast suspensions was determined and adjusted (1 OD_600_ = 3 x 10^7^ CFU/ml) [84]. Sterile Milli-Q™ ddH_2_O was used in all experiments.

### CRISPR/Cas9 mediated gene inactivation

The CRISPR/Cas9 gene editing was used to inactivate both alleles of *SSY1* (C2_04060C), *CSH3* (C4_03390W), *CAR1* (C5_04490C), *PUT1* (C5_02600W), *DUR1,2* (C1_04660W), *IRA2* (C1_12450C), *PUT1* (C5_02600W), *PUT2* (C5_04880C) or *PUT3* (C1_07020C). Sequences of synthetic guide RNAs (sgRNAs), repair templates, and verification primers are listed in Table S2. The solo system plasmids pV1093 or pV1524 were used [85, 86]. These plasmids contain a cassette comprised of the *Candida*/*Saccharomyces* codon-optimized *CAS9* endonuclease gene, *NAT* gene (recyclable in pV1524), sgRNA cloning site, and flanking sequences for genomic integration. For pV1093 and its derivative plasmids, the cassettes were integrated in one of the *ENO1* loci, whereas pV1524 and its derivatives where integrated in one of the *NEUT5* loci. The sgRNAs were designed as described [87] and were inserted in pV1093 or pV1524 by linker ligation. To summarize, oligo pairs p43/p44 (*SSY1*), p49/p50 (*CSH3*), p55/p56 (*CAR1*), p61/p62 (*DUR1,2*), p67/p68 (*IRA2*), p73/p74 (*PUT1*), p79/p80 (*PUT2*), and p85/p86 (*PUT3*), were separately phosphorylated and annealed prior to ligating them to dephosphorylated *Esp*3I (*Bsm*BI)-digested pV1093 or pV1524. Ligation reactions were purified and introduced into *E. coli* by electroporation. Transformants were selected on solid LB+Cb (or +Nou for pV1524 cloning) incubated overnight at 37 °C. Plasmids were sequenced using primer p91 (FS95). Plasmids (3 to 6 μg) containing the 20-bp sgRNA for *SSY1* (pFS013), *CSH3* (pFS017), *CAR1* (pFS024), *DUR1,2* (pFS039), *IRA2* (pFS028), *PUT1* (pFS080, pV1093 derivative), *PUT1* (pFS088, pV1524 derivative), *PUT2* (pFS083) and *PUT3* (pFS084) were digested with *Kpn*I and *Sac*I to release the cassette. Repair templates (RT) containing stop codon and specific restriction site were produced by template-less PCR using oligo pairs p45/p46 (*SSY1*), p51/p52 (*CSH3*), p57/p58 (*CAR1*), p63/p64 (*DUR1,2*), p69/p70 (*IRA2*), p75/p76 (*PUT1*), p81/p82 (*PUT2*), and p87/p88 (*PUT3*). PCR-purified digested plasmid and repair templates were co-transformed into *C. albicans* cells at a 1:3 ratio (w/w, plasmid:repair template). The hybrid lithium acetate/DTT-electroporation method, with minor modifications, was used for transforming *C. albicans* [88]. After applying 1.5 kV of electric pulse, cells were recovered in YPD medium supplemented with 1 M sorbitol for at least 4 h and then plated on YPD+Nou plates; Nou^R^ colonies were selected 2 days after plating. Nou^R^ transformants were pre-screened according to the expected phenotype prior to PCR and restriction analysis using primers and restriction enzymes indicated in Table S2 (**Fig. S3**).

### Full genome shotgun sequencing

Genomic DNA was isolated from *put1*-/- (CFG139), *put2*-/- (CFG207), *put3*-/- (CFG146), *put1*-/- *put2-/-* (CFG159) and CRISPR/Cas9 control strains CFG181 (pV1093) and CFG182 (pV1524) and sequenced. Prior to library construction, extracted DNA was purified with Agencourt AMPure^®^ XP beads (Beckman Coulter, USA) in order to remove short sequences (<100 bp). Aliquots (25 μl) of DNA were mixed with 45 μl of AMPure beads with a ratio of 1:1.8 and incubated 15 min. Initial DNA concentrations following purification were evaluated using Quant-iT PicoGreen dsDNA Assay kit (ThermoFisher, USA) as described (Logares & Feng, 2010). Absorbance was measured at 530 nm, using a Tecan Ultra 384 SpectroFluorometer (PerkinElmer, USA).

Library construction was carried out with the QIAGEN-FX kit (Qiagen, Germany) with a DNA input of 100 ng DNA per sample and a digestion time of 13 min without enhancer. Following fragmentation, adapter sequences were ligated, and ligated DNA fragments were amplified by 9 cycles of PCR and DNA was purified with AMPure^®^ XP beads. The quality of the library samples were evaluated with an Agilent Bioanalyzer using DNA1000 cartridges. The average length of the fragments excluding adapter sequences was 455 bp.

Prior to sequencing, the samples were denatured with 0.2 N NaOH. A final volume of 570 μl of pooled library was mixed with denatured Phix control (30 μl) and loaded on an Illumina Mi-Seq 2×300 flow-cell and reagent cartridge. De-multiplexing and removal of indexes and primers were done with the Illumina software v. 2.6.2.1 on the instrument according to the standard Illumina protocol. Initial de novo assembly of quality controlled reads was done with SPADES v. 3.11.1 and standard settings [89]. Mapping of assembled contigs was done with Ragout v 2.0 [90] using Sibelia for synteny detection[91]. Visualization of results and generation of reports on the assembly quality and other factors were done with QUAST v. 4.6.1 [92].

### NanoLuc transcription-translation reporter of SPS-sensor activation

The NanoLuc-PEST (Nlucp) construct was used to create the reporter of SPS-sensor dependent transcription (**Fig. S2)**. The presence of PEST sequences ensures rapid degradation of NanoLuc luciferase, thereby enhancing sensitivity [39]. Up- and downstream regions of the *CAN1* ORF were amplified using genomic DNA from PMRCA18 as template and primers p92/p93 (0.9 kB upstream) and p96/p97 (0.98 kB downstream) (**Table S2)**. An approximately 0.7 kB Nlucp gene sequence was amplified from plasmid pCA873 [93] using primers p94/p95. These amplicons were digested with appropriate FastDigest enzymes (Thermo Scientific) and purified; i.e., the *CAN1* upstream amplicon was digested with *Kpn*I/*Xho*I, the *CAN1* downstream with *Xba*I/*Not*I, and Nlucp DNA fragment with *Xho*I/*BamH*I. Using T4 DNA Ligase (Thermo Scientific), the upstream fragment was first ligated to *Kpn*I/*Xho*I-digested pSFS2a vector [88] creating pFS006. The purified Nlucp DNA was then ligated into *Xho*I/*BamH*I restricted pFS006 creating pFS007. Finally, the downstream fragment was ligated into *Xba*I/*Not*I restricted pFS007 creating pFS010. The plasmids were introduced into *E. coli* and transformants selected on LB+Cm+Nou plates incubated at 30 °C. The desired reporter construct, purified from *Kpn*I/*Not*I restricted pFS010, was introduced into *C. albicans* wildtype (PMRCA18) and SPS-deficient mutant strains (*ssy1*Δ/Δ, *ssy5*Δ/Δ, and *stp2*Δ/Δ) by electroporation. Selection was carried out on YPD+Nou and NAT^R^ clones carrying the integrated Nlucp construct were identified by PCR.

For analysis of amino acid-induced SPS-sensor activation, Nano-Glo^®^ Luciferase Assay System (Promega GmbH, Germany) was used following the manufacturer’s protocol. Briefly, log phase SD cultures were first standardized to OD ≈ 0.8 before adding 50 μl of the cell suspension into each well of Nunc 96 well microplate (white). Then, cells were induced with 50 μM of the indicated amino acids for 2 h at 30 °C. Fifty microliters (50 μl) of Nano-Glo substrate diluted 1:50 in the supplied lysis buffer was added into each well of the microplate. After 3 min, bioluminescence was captured using microplate luminometer (Orion II, Berthold Technologies GmbH & Co. KG, Germany). Luminescense reading from treated wells were deducted from wells spiked with ddH_2_O serving as uninduced control.

### Filamentation assay

Solid filamentation assay was performed as described [14]. Briefly, cells from overnight YPD liquid cultures were harvested, washed once, and resuspended in sterile ddH_2_O. The cell density of cell suspensions was adjusted to OD_600_ ≈ 8 before spotting 10 μl onto solid media. Plates were allowed to dry at room temperature before incubating at 37 °C as indicated to allow macrocolonies to form. Filamentation assays in the presence of metabolic inhibitors, nor-NOHA or L-THFA, were performed in a 6-well microplate format (~5 ml/well); otherwise, all assays were carried out using standard Petri plates (~35 ml/plate). For filamentation assays in liquid cultures, cells were washed and then adjusted to OD_600_ ≈ 25. Cells were diluted in pre-warmed liquid medium at OD_600_ ≈ 0.5 and then incubated at 37 °C with vigorous agitation for the specified time. Cell morphologies were assessed under epifluorescence microscopy using calcofluor white stain (CFW, Fluorescent Brightener 28, 1 mg/ml; Sigma).

### qRT-PCR

Hyphal specific gene (HSG) expression in 24 h old macrocolonies was analyzed as follows: using a sterile glass slide, three to four macrocolonies of wildtype strain (PMRCA18) were collected by scraping and suspended in 1 ml of ice-cold PBS. Cells were harvested by centrifugation at 10,000 x g for 3 min (4 °C), snap frozen in liquid nitrogen and then stored at - 80 °C until processed for RNA extraction. Gene expression in liquid grown cells was analyzed as follows: cells from overnight YPD cultures were harvested by centrifugation, washed and resuspended at an OD_600_ ≈ 25 in pre-warmed liquid medium and incubated at 37 °C for 2- and 4- h before harvesting the cells by centrifugation; the cell pellets were snap frozen in liquid nitrogen. For arginine catabolic gene expression analysis, SC5314 was used as wildtype strain. Briefly, cells from log phase YPD culture growing at 30 °C were harvested, washed 3X with PBS, diluted in pre-warmed YNB+Arg medium (pH = 6.0, without BCP) at an OD_600_ ≈ 0.5, and then incubated for 1 h at 37 °C under aeration. A portion of the washed cells were snap-frozen in liquid nitrogen to serve as reference (t = 0). Following 1 h incubation, cells in YNB+Arg were immediately harvested and then snap-frozen in liquid nitrogen for RNA extraction. To analyze the dependence of *GAP* genes expression to SPS pathway (i.e., Ssy1), wildtype (PMRCA18) and *ssy1*Δ/Δ (YJA64) cells were grown to log phase in SD medium at 30 °C before spiking with 1 mM of glutamine or ddH_2_O for 30 min. Cells were collected from induced (glutamine) and non-induced (ddH_2_O) cultures and snap-frozen in liquid nitrogen.

Total RNA was extracted from frozen cell pellets using RiboPure-Yeast Kit (Ambion®, Life Technologies) essentially following the instructions of the supplier with the exception that cells were subjected to extra bead-beating step (Bio-Spec; 1 × 60 sec, 4 M/s). DNase-treated RNA extracts were reverse-transcribed using SuperScript III and Random Primers (Invitrogen, Life Technologies). cDNA preparations were diluted 1/40 in ddH_2_O and 5 μl were used as template for qPCR using KAPA SYBR Green (Kapa Biosystems). Gene specific primers (500 nM) were added and reactions were performed in a Rotor-Gene 6000 (software version 1.7). The ΔΔCt method (2^-ΔΔCt^) was used to quantitate the relative levels of gene expression.

### ATP quantification

A bioluminescence-based ATP detection kit (Molecular Probes, Invitrogen) was used to quantify ATP levels in macrocolonies grown on S<u>X</u>D medium as indicated. ATP was extracted from eight, 24 h-old macrocolonies harvested using a sterile glass slide and then suspended in 1 ml sterile ice-cold Tris Buffered Saline (TBS; 50 mM Tris-Cl, pH 7.5, 150 mM NaCl). Cells were harvested at 10,000 × g for 3 min (4°C) before re-suspending the entire pellet in TCA buffer containing 100 mM Tris-HCl (pH = 8.0), 10% trichloroacetic acid (TCA), 25 mM ammonium acetate, and 4 mM EDTA. Cell suspension was transferred to pre-chilled tubes containing glass beads and then subjected to bead beating (Bio-Spec; 5 × 1 min, 4 M/s with 2 min on ice between pulses). Cell lysates were collected and a portion of the supernatant was analyzed for ATP following the instruction of the manufacturer. Luminescence was analyzed using microplate reader (Berthhold) using 1 sec integration time. A portion of the same lysate was used to determine total protein concentration using the bicinchoninic acid (BCA; Sigma) assay. Results presented are average of ATP normalized to total protein concentration analyzed from 3 biological replicates; each replicate is an average of 2-3 technical replicates.

### Immunoblotting

For Stp2 cleavage analysis, cells expressing Stp2-HA (PMRCA48) were grown to saturation in SD liquid medium overnight at 30 °C and then refreshed the following morning in 25 ml of fresh SD medium at a starting OD_600_ ≈ 0.3. Cells were grown in a 30 °C-shaker to an OD_600_ of 1.5-2.2. For induction experiments, a 500-μl aliquot of log phase culture were separately added to tubes containing the indicated amount and type of amino acids or an equal volume of water for control, and then incubated for 5 min at 30 °C in a thermoblock shaking at 700 rpm. For Put2-HA expression analysis, cells from overnight YPD cultures were harvested, washed and then grown as indicated. Whole cell lysates were prepared using NaOH/TCA method as described previously with minor modifications [94]. Cells were lysed on ice with 280 μl of ice-cold 1.85 M NaOH with 7% ß-Mercaptoethanol for 15 min; proteins were precipitated ON at 4 °C by adding the same volume of cold 50% TCA. Protein pellets were quickly washed with ice-cold 1 M Tris base (pH = 11) and then resuspended in equal volume of Tris-HCl (pH = 8.0). In some instances, as indicated, due to highly variability in protein recovered from certain types of cells (i.e., yeast and filamentous forms) sample loading was normalized based on protein content. Samples were denatured in 2X SDS sample buffer at 95-100 °C for 5 min, the proteins were resolved in sodium dodecyl sulfate-polyacrylamide gel electrophoresis (SDS-PAGE) using 4-12% pre-cast gels (Invitrogen) and analyzed by immunoblotting on nitrocellulose membrane according to standard procedure. For Stp2-HA and Put-HA detection, HRP-conjugated anti-HA antibody (Pierce) was used at 1:2,500 dilution. For loading control, HRP-conjugated rat monoclonal α-tubulin antibody [YOL1/34] (Abcam) was used at 1:10,000 dilution. Membranes were blocked using TBST (TBS + 0.1% Tween) containing 10% skimmed milk; antibodies were diluted in TBST containing 5% skimmed milk. Immunoreactive bands were visualized by enhanced chemiluminescent detection system (SuperSignal Dura West Extended Duration Substrate; Pierce) using ChemiDoc MP system (BioRad). Densitometric analyses were performed using ImageJ.

### Active Ras1-Pull Down Assay

Active Ras1 (Ras1-GTP) was analyzed in macrocolonies using Pierce Active Ras Pull-Down Kit (Thermo Scientific) following the manufacturer’s instructions, but with an extra bead-beating step to ensure optimal disruption of cells. Five 24 h-old macrocolonies were scraped, pooled, and suspended in 1 ml ice-cold TBS in 2-ml microcentrifuge tubes (with caps). Cells were collected by centrifugation at 10,000 x g for 3 min (4 °C) and then resuspended in 400 μl of Lysis/Binding/Washing buffer (1X, Pierce kit) supplemented with protease cocktail (cØmplete™ mini, EDTA-free; Roche) and 1 mM PMSF. Pre-chilled glass beads were added, cell suspensions were subjected to multiple cycles of bead beating (6 × 40 sec, 4 M/s, 2 min on ice between pulses). After an initial clarification step at 1,000 rpm for 5 min, supernatants were collected and total protein was determined using the BCA assay. The concentration of protein in lysates was adjusted to 2 mg/ml using the lysis buffer as diluent and then 500 μg of protein was used for the immunoprecipitation. We used 12.5 μg protein for input and eluted bound protein in 25 μl. Proteins were resolved by SDS-PAGE and analyzed by immunoblotting. Total Ras and active Ras-GTP were probed with primary monoclonal anti-Ras clone X (1:300) included in the kit, and secondary goat anti-mouse antibody (1:10,000; Pierce). For loading control, α-tubulin conjugated to HRP (1:10,000) was used. Membranes were blocked and the primary antibody diluted in TBST containing 3% BSA; the secondary and loading control antibodies were diluted in TBST containing 5% skimmed milk. Results presented are representative of at least 3 independent experiments.

### Growth Assays

For drop plates, cells from log phase YPD cultures grown at 30 °C were harvested, washed, and then adjusted to OD_600_ ≈ 1. Five microliters of 10-fold serially diluted cell suspension were spotted onto the surface of the indicated S<u>X</u>D media and incubated at 30 °C for 2-3 days and photographed. For liquid assays, washed cells from log phase YPD cultures were diluted in the indicated S<u>X</u>D liquid medium to a starting OD_600_ ≈ 0.05, and 300 μl were transferred into each well of a 10 x 10-well microplate and grown continuously for > 20 h at 30 °C with constant agitation. OD_600_ readouts were captured using BioScreen C MBR analyzer (Oy Growth Curves Ab Ltd, Helsinki, Finland).

### Resazurin Reduction Assay

The membrane permeant, non-destructive redox indicator, Resazurin (Sigma), was used to measure the metabolic activity of intact cells growing in S<u>X</u>D. Briefly, cells from overnight YPD cultures were harvested, washed once with sterile ddH_2_O, and adjusted to OD_600_ ≈ 0.01 (~3 x 105 CFU/ml) using the YNB-glucose base medium (~1.05x strength, pH = 6.0). Using a multi-channel pipette, 95 μl of this cell suspension were added to the well of a 96-well microplate followed by addition of 5 μl of 200 mM amino acid stock (10 mM final concentration). Plates were incubated at 37°C for 2 h with agitation protected from light. After 2 h, 20 μl of filtered Resazurin dye (0.15 mg/ml) was added to each well and incubated for 2 h at 37 °C before measuring the fluorescence (560 nm excitation/590 nm emission) using EnSpire microplate reader (PerkinElmer).

### TTC Overlay Assay

Macrocolonies grown on the indicated plates for 24 h were overlaid with 2 ml of molten TTC-agar solution (50-55 °C) containing 0.1% TTC (2,3,5 triphenyltetrazolium chloride; Sigma) dissolved in 6.7 mM potassium phosphate buffer (PPB, pH = 7.0) with 1% agar [15]. Plates were photographed 30 min after the overlaid solution became solid.

### Extracellular oxygen consumption assay

Oxygen consumption assay was performed in *C. albicans* grown in synthetic proline medium containing the indicated carbon source (i.e., 2% glucose (SP<u>D)</u>, 0.2% (SP<u>D_0.2%_</u>), or 1% glycerol (SP<u>G)</u> using the Extracellular Oxygen Consumption Assay (Abcam, ab197243) following manufacturer’s protocol. Briefly, cells from log phase YPD culture were harvested, washed 3X with PBS, and then diluted in the indicated media at OD_600_ ≈ 0.3. A 150 μl cell suspension was added into each well of a 96-well microplate with black walls and clear bottom. Ten microliters (10 μl) of Extracellular Oxygen Consumption Reagent or medium were then added into each well, mixed gently by moving the plate on a circular motion, and then spiked with either medium or control. FCCP (final conc. 10 μM) and antimycin (final conc. 10 μg/ml) were used as positive and negative controls, respectively. Plates were analyzed using Enspire microplate reader using Time Resolved Fluorescence (TRF). Signals were read every 90 sec for 120 repeats with optimal delay time of 30 μs and gate (integration) time of 100 μs. Signal from wells without cells were used as background signal.

### Quantification of proline

The concentration of proline in media and in cell extracts was analyzed using the quantitative ninhydrin method [95]. Proline utilization was assessed as follows: cells grown overnight in YPD were washed and resuspended to an OD_600_ ≈ 0.5 in pre-warmed synthetic proline media containing 10 mM of proline and the indicated carbon source. The cultures were incubated under constant aeration for 2 h at 37 °C, and the amount of proline in culture supernatants was analyzed. Proline utilization was defined by comparison to non-inoculated media.

### *C. albicans* co-culture with murine macrophages

The murine macrophage cell line RAW264.7 (ATCC) was cultured and passaged in Dulbecco’s modified Eagle’s medium/high glucose (HyClone, GE Healthcare Life Sciences, Amersham, UK) supplemented with 10% fetal bovine serum, 100 U/ml penicillin and 100 μg/ml streptomycin (hereafter referred as D10) at 37°C with 5% CO_2_. Prior to co-culture with *C. albicans*, RAW264.7 cells (1 × 10^6^) in D10 medium were seeded on a 24-well microplate containing sterile cover slips and were allowed to adhere overnight in a humidified chamber at 37°C and 5% CO_2_. Fungal cells (3 × 10^8^) were harvested from overnight YPD cultures and stained with 1 mg/ml FITC solution in 0.1 M NaHCO_3_ buffer (pH = 9.0) in the dark for 15 min at 30 °C. Cells were washed 3X with PBS before resuspending in equal volume of PBS. Fungal cells were added to macrophage at MOI of 3:1 (Candida:Macrophage, C:M) and were then allowed to interact for 30 min. Non-phagocytosed cells were removed by washing the cells at least 5X with pre-warmed Hank’s Balanced Salt Solution (HBSS) and 1X with D10 medium. Cells were allowed to interact for an additional 4 h in fresh D10 medium before fixing with 3.7% formaldehyde-PBS for 15 min in the dark at room temperature. Fixed cells were then washed 3X with PBS before staining with calcofluor white (10 μg/ml) for 1 min. After 2X PBS washing, coverslips were mounted on glass slides using ProLongTM Gold antifade reagent (Invitrogen). Images were obtained using LSM 800, 63x/1.2 oil.

### *C. albicans* killing by murine macrophages

The survival of *C. albicans* co-cultured with macrophages was assessed by colony forming units (CFU) analysis. Briefly, RAW264.7 cells in D10 were seeded into a 96-well microplate at a density of 1 x 10^5^ per 200 μl and allowed to adhere overnight. *C. albicans* cells from overnight YPD cultures were processed without staining and added at a MOI of 3:1 (C:M). The co-cultures were incubated for 3 h prior to assessing fungal cell viability by CFU; each well was treated to final concentration of 0.1 % Triton X-100 for 2 min to lyse macrophage and serial dilutions were prepared and plated onto YPD. CFUs were counted 2 days after incubation at 30 °C. The ability of macrophages to kill *C. albicans* (% killing) was determined by comparison of fungal CFU recovered in the absence of macrophages.

### Indirect immunofluorescence microscopy of phagocytosed *C. albicans*

RAW264.7 cells were co-cultured with *C. albicans* cells, CFG185 (*PUT2/PUT2-HA*), for 90 min on glass coverslips at a MOI of 5:1 (C:M). Cells were fixed in 3.7% formaldehyde-PBS for 15 min, and permeabilized in 0.25% Tween-20 for 15 min, both incubations were at room temperature. The fixed and permeabilized cells were incubated in zymolyase buffer (2U zymolyase 100T (Zymo Research, Irvine, CA, USA), 10 mM DTT in PBS) for 1 h at 30 °C. After washing, cells were incubated at room temperature in 0.25% Tween-20 for 10 min and blocked in 5% FBS for 30 min. Cells were incubated overnight at 4 °C with rat anti-HA (Roche, Germany, #1867423) and rabbit anti-Lamp1 (Abcam, UK, #ab24170) primary antibodies diluted 1:500 in 0.25% Tween-20. Cells were washed with PBS and incubated 2 h with Alexa flour 488 goat anti-rabbit (Invitrogen, Eugene, OR, USA #A11034) and Alexa flour 555 goat anti-rat (Invitrogen, Eugene, OR, USA #A11034) secondary antibodies diluted 1:500 in 0.25% Tween-20. Images were captured on a Zeiss 510 Meta confocal microscope, 63x/1.4 oil. Orthogonal views were constructed in FIJI imaging software.

## Acknowledgments

The authors would like to thank the members of Claes Andréasson, Martin Ott, and Per Ljungdahl laboratories (SU), and members of the Marie Curie-ITN ImResFun program for constructive comments throughout the course of this work. Gratitude is extended to Valmik Vyas and Gerard Fink (MIT, Cambridge, MA, USA) for providing the CRISPR/Cas9 cassettes and for the valuable suggestions. Patrick van Dijck (KUL, Leuvan, Belgium), Karl Kuchler (MUV, Vienna, Austria), Kenneth Nickerson and Ruvini Pathirana (UNL, Lincoln, NE, USA) are gratefully acknowledged for supplying strains and discussions; and Nora Grahl and Deborah Hogan (Dartmouth Medical School, Hanover, NH, USA) for helpful discussions. We also thank Stina Höglund, the Imaging Facility-Stockholm University, for assistance in microscopy. This work was supported by EU grant MC-ITN-606786 (ImResFun) and Swedish Research Council VR-2015-04202 (POL).

## Supporting Information

**Fig S1. Hyphae-specific gene (*HSG*) expression in *C. albicans* grown in the presence of inducing amino acids.**

**Fig S2. NanoLuc™ luciferase assay for analysis of Stp2 target gene expression.**

**Fig S3. CRISPR/Cas9-mediated gene inactivation in *C. albicans*.**

**Fig S4. Metabolic activity of *C. albicans* grown in the presence of inducing and non-inducing amino acids.**

**Fig S5. Growth curves of arginase-pathway mutants in different nitrogen sources.**

**Fig S6. Neutralization of medium containing amino acid as sole carbon and nitrogen source remains intact in mutant lacking *DUR1,2*.**

**Fig S7. Carbon source and mitochondria-dependent oxygen consumption of *C. albicans***.

**Table S1. Strains used in this study.**

**Table S2. Primers used in this study**

